# Brain endothelial STING1 activation by *Plasmodium*-sequestered heme promotes cerebral malaria via type I IFN response

**DOI:** 10.1101/2022.02.14.480268

**Authors:** Teresa F. Pais, Hajrabibi Ali, Joana Moreira da Silva, Nádia Duarte, Rita Neres, Chintan Chhatbar, Rita C. Acúrcio, Rita C. Guedes, Maria Carolina Strano Moraes, Bruno Costa Silva, Ulrich Kalinke, Carlos Penha-Gonçalves

## Abstract

Cerebral malaria (CM) is a life-threatening form of *Plasmodium falciparum* infection caused by brain inflammation. Brain endothelium dysfunction is a hallmark of CM pathology, which is also associated with the activation of the type I interferon (IFN) inflammatory pathway. The molecular triggers and sensors eliciting brain type I IFN cellular responses during CM remain largely unknown. We herein identified the stimulator of interferon response cGAMP interactor 1 (STING1) as the key innate immune sensor that induces *Ifnβ1* transcription in the brain of mice infected with *Plasmodium berghei* (*Pba*). This STING1/IFNβ-mediated response increases brain CXCL10 governing the extent of brain leucocyte infiltration and blood-brain barrier (BBB) breakdown, and determining CM lethality. The critical role of brain endothelial cells (BECs) in fueling type I IFN-driven brain inflammation was demonstrated in brain endothelial-specific IFNβ-reporter and STING1-deficient *Pba*-infected mice, which are significantly protected from CM lethality. Moreover, extracellular particles (EPs) released from *Pba*-infected erythrocytes activated STING1-dependent type I IFN response in BECs, a response requiring intracellular acidification. Fractionation of the EPs enabled us to identify a defined fraction carrying hemoglobin degradation remnants that activates STING1/IFNβ in the brain endothelium, a process correlated with heme content. Notably, stimulation of STING1-deficient BECs with heme, docking experiments and *in vitro* binding assays unveiled that heme is a putative STING1 ligand. This work shows that heme resultant from the parasite heterotrophic activity operates as an alarmin triggering brain endothelial inflammatory responses via STING1/IFNβ/CXCL10 axis crucial to CM pathogenesis and lethality.

**Significance:** CM results from loss of blood-brain endothelial barrier function caused by unrestrained inflammatory response in the natural course of infection by *Plasmodium* parasites. However, the role of brain endothelium in triggering inflammatory mechanisms is still undetermined. We found that the innate immune sensor STING1 is crucial for production of IFNβ in brain endothelial cells in *Plasmodium*-infected mice. This in turn stimulates CXCL10-mediated recruitment of leukocytes and subsequent brain inflammation and tissue damage. We identified within extracellular particles released from *Plasmodium*-infected erythrocytes, a fraction containing products of hemoglobin degradation, namely heme, which we show can bind STING1. Our results unravel a new angle of CM pathogenesis: heme contained in particles triggers the STING/IFNβ/CXCL10 axis in brain endothelial cells.

## Introduction

Malaria, the disease caused by Plasmodium infection, accounted for over 400 thousand deaths worldwide in 2019 (World Malaria Report 2019). Despite the available malarial treatments, children under 5 years of age can develop cerebral malaria (CM) upon infection with *Plasmodium falciparum*, facing the highest risk of severe malaria morbidity and mortality (15-20%) (1). Brain swelling with increased compression of respiration centers in the brain stem predicts a fatal CM outcome (2). Children who survive CM are often affected by long-term neurological sequelae (1).

Infected erythrocytes (IE), parasite components released during the blood-stage infection (e.g. malarial hemozoin and labile heme), and systemic inflammatory factors (eg. CXCL10, TNF, IL1β and IFNγ) elicit an unfettered brain endothelial cell response that compromises regular endothelium function (reviewed in (3)). In particular, BBB integrity is lost, which results in leakage of plasma proteins and water into the cerebral parenchyma and consequent intracranial hypertension (reviewed in (4)). However, whether brain endothelial cells (BECs) are bystander activated by systemic inflammatory factors or the actual source of brain inflammation and pathology remains largely unexplored in CM. Over the last years, the type I interferons, IFNα and IFNβ, emerged as key mediators in innate immune responses to malaria, promoting brain immunopathology (reviewed in (5)). In children with uncomplicated malaria, activation of the type I IFN response was associated with protective immunity (6). However, in children with CM, polymorphism in regulatory regions of type I IFN receptor (IFNAR1 subunit) gene were associated with increased type I IFN signaling and CM development (7). In the experimental mouse model of CM, despite the protective role of type I IFN in early infection (8, 9), several reports show that ablation of IFNAR1 signaling prevents BBB disruption and pathogenesis of experimental CM driven by parasite-specific cytotoxic CD8^+^ T cells (10, 11). In addition, mice deficient in type I IFN signaling molecules—like transcription factors interferon regulatory factor 3 (IRF3) and 7 (IRF7) and the upstream TAK-binding kinase 1 (TBK1)—are protected from CM (12). Moreover, a point mutation in the ubiquitin-specific protease 15 (USP 15) deters the type I IFN response in the mouse brain, which dampens brain inflammation and increases survival to CM (13).

Several pattern recognition receptors (PRRs) underlie inflammatory responses during *Plasmodium* infection by binding to *Plasmodium*-derived components (14), such as GPI anchors, plasmodial DNA and RNA, and hemozoin—a byproduct of hemoglobin digested by the parasite, consisting of crystals made of polymerized heme (15). Labile heme, a byproduct of hemolysis during *Plasmodium* infection, can also signal via PRR (16) and contribute to the pathogenesis of CM (17, 18).

The cGAS-STING1 pathway is a major sensor of cytosolic DNA that can recognize AT-rich motifs in plasmodial DNA and induce IFNβ transcription in human monocytes and mouse macrophages (12, 19, 20). Interestingly, extracellular vesicles derived from *Plasmodium*-IE (19) and complexes of plasmodial DNA with hemozoin (12) were proposed as carriers of parasite DNA into the cytosol of human monocytes.

BECs can also capture and cross-present malarial antigens to parasite-specific CD8^+^ T cells (21). Two mechanisms by which malarial antigens could reach the endothelial cell endosomal and cytosolic compartments were proposed, one through membrane fusion with infected-erythrocytes, and another by uptake of merozoites and digestive vacuoles (21, 22).

Ubiquitous expression of type I IFN and IFNAR1 poses a significant challenge to the identification of specific cellular and molecular contributors to type I IFN signaling activation in CM development. Although BECs are within the first cells to sense the IE, their role in type I IFN-mediated innate immune responses during CM remains elusive. Here, we demonstrate that in the brain, specifically in BECs, STING1 activation by parasite-sequestered heme present in particles released from Plasmodium-IE is a critical mechanism underlying CM pathogenesis.

## Results

### Brain IFNβ production via STING 1 activation correlates with CM

There is indirect evidence connecting brain type I IFN responses with tissue pathology associated with CM in *Pba*-infected mice (23). We used an IFNβ-luciferase reporter mouse line (24) to directly test whether IFNβ induction in the brain was triggered by *Pba* infection. We detected strong luminescence (fig. S1A) and significantly increased luciferase activity in brain tissue as well as in the lung of mice developing CM (Fig. 1A). In contrast, mice infected with *PbNK65,* a parasite strain that does not cause CM, induced weak or no luminescence signal (fig. S1B). The IFNβ induction in the brain correlated with a type I response spread in three main brain regions—olfactory bulb, cerebrum and cerebellum—as indicated by the expression of four type I IFN response interferon-stimulated genes (*Irf1*, *Irf7*, *Ifit1* and *Cxcl10*) (fig. S1C). We then assessed the contribution of IFNβ to CM pathogenesis. BBB integrity was measured by EB quantification in the brain after perfusion at day 6 post infection (PI). BBB leakage of EB was lower in the brains of IFNβ KO than in the brains of wild type (WT) mice (Fig. 1B and fig. S1D). Accordingly, CM was less lethal in IFNβ KO mice than in WT controls (Fig 1D). This decrease in lethality was not attributable to differences in parasite burden between IFNβ KO and WT mice, and was not affected by sex (fig. S1E).

**Fig. 1.**
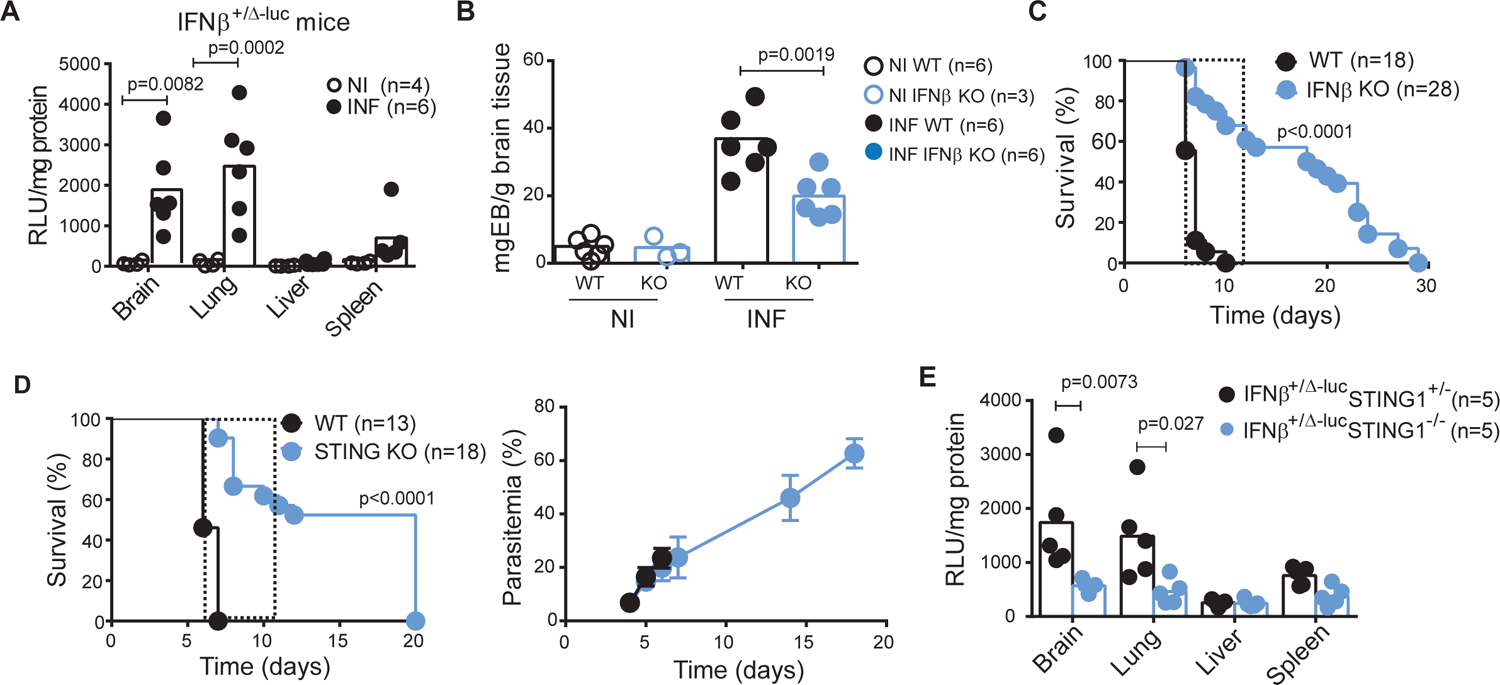
Cerebral malaria pathogenesis is associated to type I IFN responses induced by STING1 activation in *Plasmodium* infection. **(A)** *Ifnβ1* gene expression in the indicated organs of IFNβ-Luc reporter (IFNβ^+/Δβ-luc^) mice infected with *Pba* (day 6 PI) or non-infected was ascertained by *ex vivo* luciferase activity normalized for tissue protein contents. The present data are representative of three independent experiments. **(B)** Blood-brain barrier integrity was evaluated by Evans Blue (EB) quantification in the brain after mouse perfusion. Data were normalized for brain weight in WT and IFNβ KO mice at day 6 PI. **(C)** Survival curve of WT and IFNβ KO mice infected with 10^6^ *Pba*-IE. **(D)** Survival and blood parasitemia curves in mice deficient for STING1. **(E)** Induction of *Ifnβ1* gene expression in the indicated organs of IFNβ-Luc reporter (IFNβ^+/Δβ-luc^STING1^+/-^) and in IFNβ-Luc reporter lacking STING (IFNβ^+/Δβ-luc^STING1^-/-^) mice infected with *Pba* (day 6 PI) was ascertained by *ex vivo* luciferase activity normalized for tissue protein amounts. Dashed line in **(C)** and **(D)** delimits the time-window of death by CM. Data points in **(A), (B)** and **(E)** represent individual mice. Data in **(A)**, **(B)** and **(E)** are analyzed by one-way ANOVA, Tukey’s multiple comparisons test. Data in **(C)** and **(D)** comprise two independent infection experiments and are compared between mouse strains with Gehan-Breslow-Wilcoxon test.

Several PRRs sense *Plasmodium* infection and initiate downstream signaling converging at the phosphorylation and activation of type I interferon regulatory family of transcription factors (IRFs)(14). Pba-IE, *Plasmodium* AT-rich DNA motifs and parasite DNA in EPs that are released from *P. falciparum*-IE were shown to activate the STING1 pathway of type I IFN induction in mouse macrophages and human monocytes in culture (12, 19). However, the role of STING1 activation in the development of CM has not been addressed. We found that *Sting1* deletion significantly delayed death by CM and conferred approximately 50% protection against CM lethality without affecting the parasitemia (Fig. 1D). Although we cannot exclude minor contributions of the other PRRs, we did not detect a significant difference in CM lethality when STING1 KO mice were compared with quadruple KO mice for *Myd88*, *Trif*, *Mavs* and *Sting1* (MyTrMaSt KO), which are impaired in all known type I response induction pathways (fig. 1F). Lower levels of IFNβ induction, measured by luciferase activity, were also found in the brains and lungs of IFNβ-reporter mice deficient for STING1 with CM (Fig. 1E). These results indicate that STING1 is a major innate immune sensor of *Pba* infection ensuing IFNβ production in the brain and subsequent BBB disruption and CM development.

### STING1 activation induces *Cxcl10* gene expression and leukocyte recruitment to the brain

In *Pba*-infected mice, brain innate immune responses associated occur before leukocyte recruitment to this organ (13, 25). We used IFNβ-reporter mice in both alleles to measure induction of type I IFN at early stages of infection. We detected a significant increase of luciferase activity in the brain of mice as early as day 4 PI (Fig. 2A). Next, we tested whether IFNβ induction by STING1 in the brain could drive expression of *Cxcl10*, a known mediator of leukocyte recruitment downstream of IFN signaling. Deletion of *Ifnβ1* or *Sting1* significantly decreased the levels of *Cxcl10* mRNA in the brain, at day 4 PI (Fig. 2B).

**Fig. 2.**
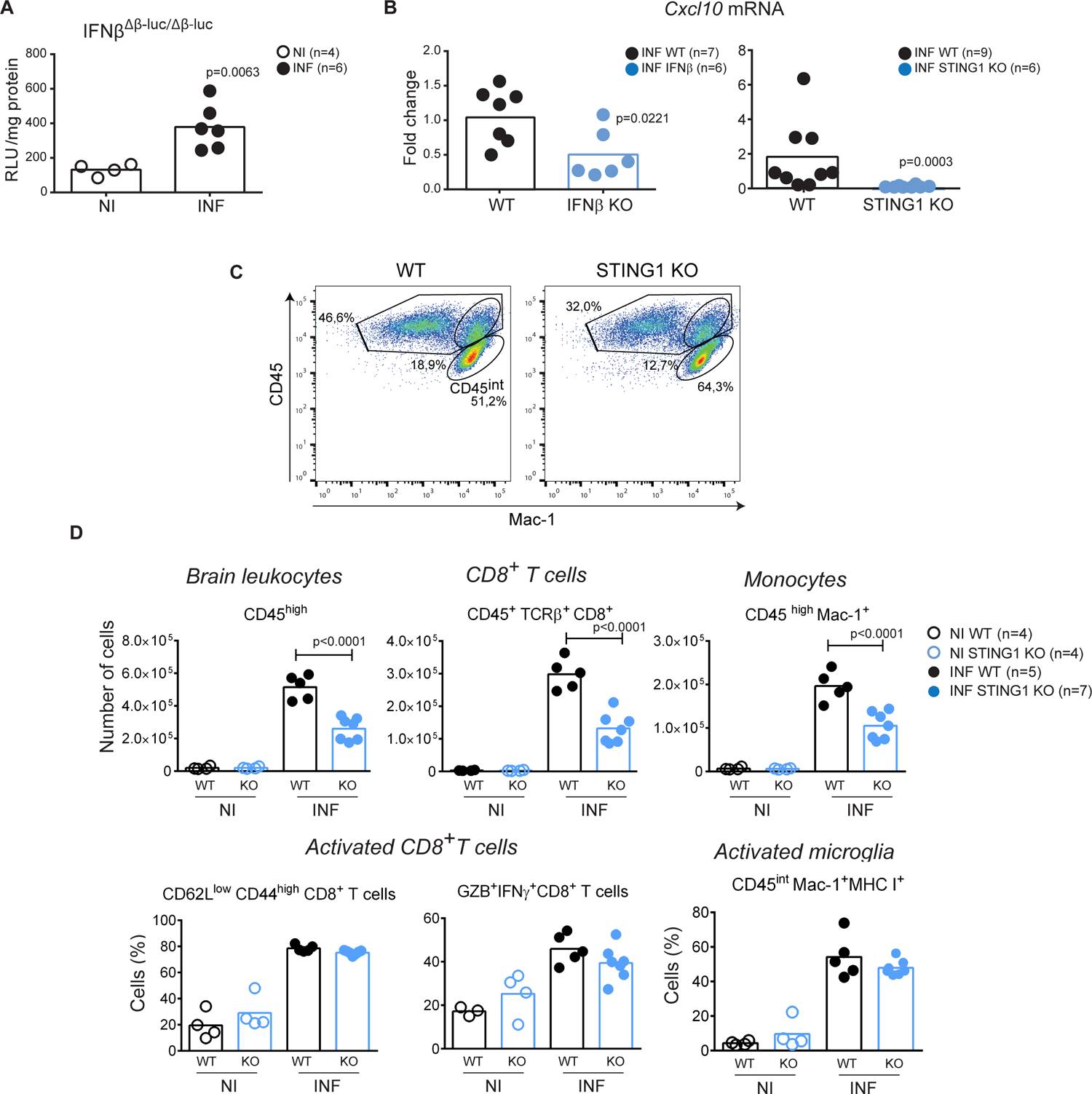
Leukocyte brain infiltration is mediated by STING1 induction of *Ifnβ* and *Cxcl10* in the brain. **(A)** *Ifnβ1* gene expression in the brain of mice with two reporter alleles for *Ifnβ1* (IFNβ^Δβ-luc/Δβ-luc^) at day 4 of infection was ascertained by *ex vivo* luciferase activity normalized for tissue protein amounts. **(B)** *Cxcl10* gene expression (qPCR) in the brain of WT and IFNβ and STING1 KO mice at day 4 of infection. **(C)** Representative flow cytometry dot blots showing the percentage of CD45^high^ cells within brain infiltrates of WT and STING1 KO mice. **(D)** Flow cytometry analysis of brain leukocytes isolated from non-infected or *Pba*-infected (day 6 PI) WT and STING1 KO mice. Extracellular staining against CD45 in combination with staining for TCRβ, CD8 and Mac-1 identified CD8^+^T cells (CD45^+^ TCRβ^+^ CD8^+^), monocytes (CD45^high^ Mac-1^+^), and microglia (CD45^int^Mac-1^+^). Activated CD8^+^T cells were identified by extracellular expression of CD62L and CD44 and intracellular staining for Granzyme B and IFNγ. MHC I expression identified activated microglial cells. Experimental groups were compared using one-way ANOVA with Tukey’s multiple comparisons test. Data points represent individual animals in **(A)**, **(B)** and **(D)**. Data are analyzed by unpaired, two-tailed *t*-test in **(A)** and **(B)**.

*Plasmodium* infection induces CXCL10 in the brain, which in turn recruits CXCR3-expressing antigen-specific pathogenic CD8^+^ T cells (26–28). We analyzed whether brain leukocyte infiltration was impaired in IFNβ KO and STING1 KO mice and found lower percentage (Fig. 2C) and numbers of leukocytes (CD45^high^), namely CD8^+^ T cells (CD45^+^TCRβ^+^CD8^+^) and monocytes (CD45^high^ Mac-1^+^), in the brains of STING1 KO mice (Fig. 2D) and of IFNβ KO mice (fig. S2A) than in those of WT mice. However, the expression of CD8^+^ T cell surface activation markers (CD44^high^/ CD62L^low^) and of Granzyme B and IFNγ was similar in WT, STING1 KO (Fig. 2D) and IFNβ KO mice (fig. S2A), which indicates that CD8^+^ T cell activation was equally effective. The low numbers of CD8^+^ T cells in the brain of infected STING1 KO mice could not be attributed to decreased local proliferation (25) as the frequency of BrdU^+^ cells amongst CD8^+^ T cells was 44.5% ± 7.74 in STING1 KO mice similar to the 33.7% ± 6.95 found in WT mice. Likewise, microglia activation assessed by MHC class I (25) expression was similar in WT and KO mice (Fig. 2D and fig. S2A).

Notably, the decrease in the number of brain-infiltrating leukocytes in the brain was not related to a decrease in the activation of splenic T cells or monocytes in STING1 KO or IFNβ KO mice (fig. S2B). This suggests that innate immune activation of STING1 leading to IFNβ production and *Cxcl10* gene up-regulation in the brain contributes to leukocyte recruitment into the brain and immunopathology in CM.

### STING1 activation and IFNβ production by brain endothelial cells contributes to CM

It is unclear whether specific brain cell types are responsible for the induction of IFNβ response to blood-borne parasites. We investigated IFNβ production in different brain cells during CM by using cell type-specific IFNβ-reporter mice (IFNβ^floxβ-luc/floxβ-luc^ mice bred to mice with Cre expression driven by cell-type specific promoters) (29). In these mice one *Ifnβ1* allele is kept wild-type while the other allele drives luciferase activity by the *Ifnβ1* floxed allele in cells that undergo Cre recombination. We found no significant induction of luciferase activity in astrocytes (GFAPCre^+^ mice) upon CM induction while a clear response was obtained 24 hours after Poly I:C injection (Fig. 3A). On the other hand, LysMCre^+^ brain cells, including different types of myeloid cells (25), showed *Ifnβ1* transcription activation upon infection as detected by increased luciferase activity in the brain (Fig. 3B).

**Fig 3.**
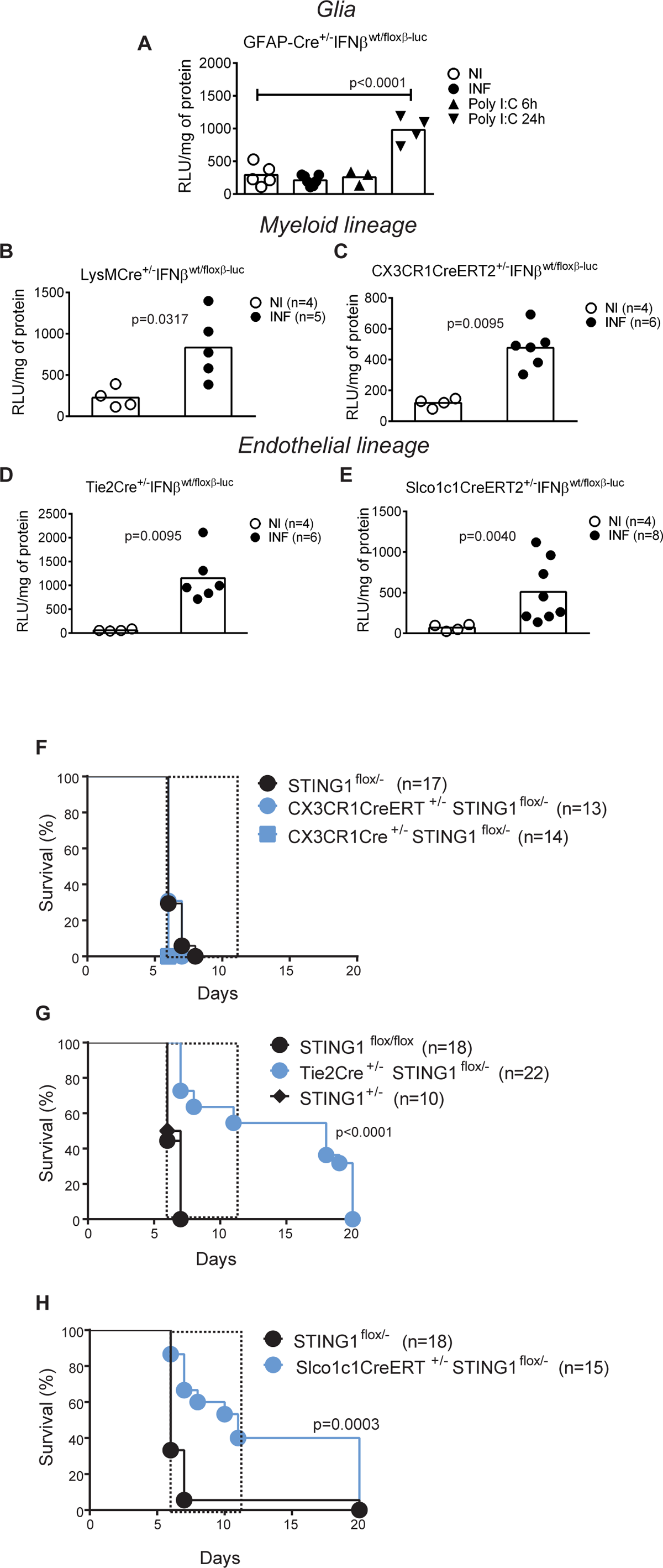
STING1 expression on brain endothelial cells is a determinant of IFNβ induction and CM lethality. *Ifnβ1* gene expression was ascertained by *ex vivo* luciferase activity in mice infected with *Pba* (day 6 PI) and non-infected mice carrying a IFNβ-Luc reporter conditionally activated in cells expressing **(A)** GFAP (glial cells), **(B)** LysM (myeloid cells), **(C)** CX3CR1 (microglia cells), **(D)** Tie-2 (endothelial and hematopoietic cells) or **(E)** Slco1c1 (brain endothelial cells). I.p. injection of Poly I:C was used as positive response control of *Ifnβ1* gene induction. Results comprise two independent infection experiments and data are analyzed with unpaired, two-tailed *t*-test. Survival curves after *Plasmodium* infection in mice carrying conditional or induced STING deletion driven by **(F)** CX3CR1Cre and CX3CR1CreERT (monocytes/microglia), **(G)** Tie2Cre (endothelial cells) or **(H)** Slco1c1CreERT (brain endothelial). Results represent at least three independent infection experiments for each mouse strain. Survival curves were compared by Gehan-Breslow-Wilcoxon test.

To determine whether IFNβ is induced in microglia, the brain resident macrophage population, during CM, we used CX3CR1CreERT2^+/-^ mice (30). After Cre induction with tamoxifen, the monocyte compartment will be replenished with bone-marrow derived cells without flox excision, while the self-renewing microglia will maintain the *Ifnβ1*-luc reporter gene active and give a luciferase signal if triggered. Under these conditions, we detected luciferase activity indicating *Ifnβ1* gene transcription in the microglial cell compartment during CM (Fig. 3C).

Induction of *Ifnβ1* in the endothelial compartment was assessed in reporter mice expressing Cre in endothelial and hematopietic cells under the Tie2 promoter (Fig. 3D), and in reporter mice carrying inducible Cre under the Slco1c1 promoter activated specifically in brain endothelial/ependymal cells (31) (Fig. 3E). Induction of luciferase activity in both mouse lines showed that brain endothelial cells contribute to local brain production of IFNβ in the brain during CM.

To determine the impact of cell-specific expression of STING1 or IFNβ in CM development, we specifically deleted these genes in different cell types by using cell-lineage specific Cre expression in mice carrying two floxed alleles or one inactivated allele and one floxed allele for the target gene. We found that absence of STING1 or IFNβ in LysM^+^ or CX3CR1^+^ cells did not prevent death by CM (Fig. 3F and fig. S3A). In contrast, ablation of STING1 or IFNβ in endothelial and hematopoietic cells in mice expressing Cre under Tie2 promoter (Fig. 3G and fig. S3B) or ablation of STING1 specifically in brain endothelial cells in Slco1c1CreERT^+/-^ mice significantly protected from CM lethality (Fig. 3H). In addition health/neurologic score on day 6 PI was improved (fig. S3C) while parasitemia was unaffected (fig. S3D). Furthermore, mice with Tie2^+^-induced STING1 deletion that survived CM showed clinical improvement after day 8 PI (fig. S3E), while parasite burden was not reduced (fig. S3F). Overall, these results unveil that STING1-dependent activation of IFNβ production in brain endothelial cells starting early in infection is a critical determinant of brain immunopathology during CM development.

### IE-derived EPs induce IFNβ via STING1 after intracellular acidification in brain endothelial cells

The generation of extracellular particles (100-1000 nm) by erythrocytes has been associated with the pathogenesis of human and rodent CM (32, 33). Also, vesicles released from *P. falciparum*-IE activate expression of pro-inflammatory cytokines (i.e TNF and IFNβ) in innate immune cells (19, 34). We asked whether extracellular particles (EPs) obtained from the serum of *P. berghei*-infected mice or from IE cell cultures can induce type I IFN signaling in primary cultures of brain endothelial cells (BECs).

Nanoparticle tracking analysis of EPs collected from mouse serum showed significantly increased mean particle size in *Pba*-infected, CM-sick mice than in non-infected mice (Fig. 4A and fig. S4A). Negative-stain transmission electron microscopy (TEM) (Fig. 4B) confirmed the presence of large particles with more than 200 nm in the serum of infected mice, while EPs isolated from non-infected animals were around 100 nm, the exosome size range (Fig. 4A). Moreover, the concentration of particles within the range of 100.5-265.5 nm increased approximately 3-fold in the blood of infected mice compared to non-infected controls (fig. 4B). The particles isolated from blood of *Pba*-infected mice that differ from those found in non-infected mice were mostly from leukocyte (CD45^+^) and erythrocytic origin (TER119^+^) as revealed by antibody staining for cell-type surface markers (Fig. 4C).

**Fig. 4.**
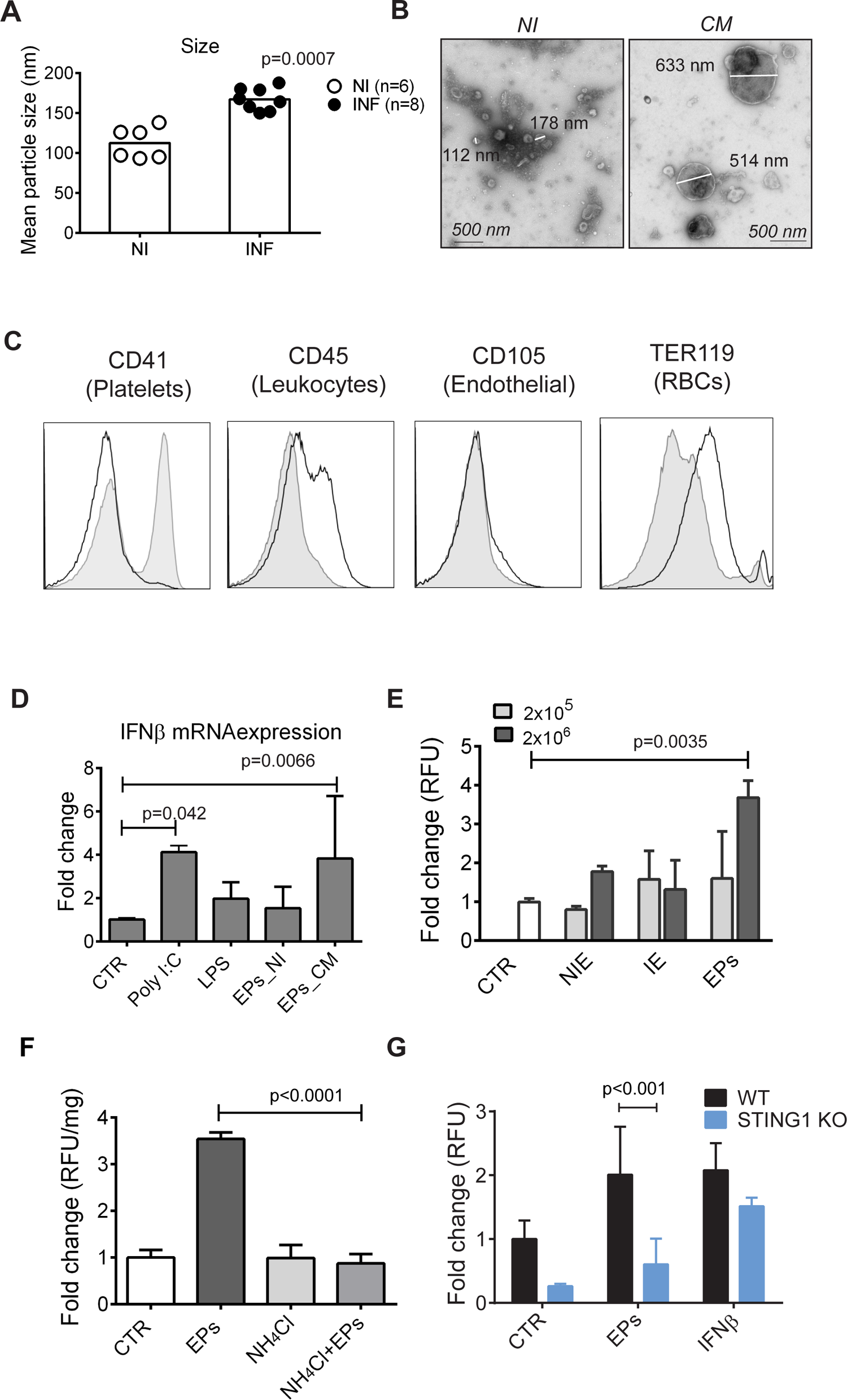
EPs isolated from blood and IE induce IFNβ in brain endothelial cells after acidic processing and in a STING1-dependent manner. **(A)** NanoSight analysis of mean particle size after serum centrifugation (20 000g) of symptomatic *Pba*-infected mice (day 6 PI) and non-infected control mice. Results represent two independent infection experiments. Non-infected mice were compared with *Pba*-infected mice by unpaired, two-tailed *t*-test. **(B)** TEM image of negative stained EPs obtained from the serum of mice with CM or non-infected animals. **(C)** Representative flow cytometry histograms of serum EPs pooled from 3-5 animals and stained with the indicated cell lineage membrane markers. EPs were collected after 20 000g centrifugation from non-infected (full histogram) and infected (empty histogram) mice. Data shown are representative of three independent experiments. **(D)** *Ifnβ1* RNA quantification in BECs stimulated with MPs isolated from the serum (600 μl) of symptomatic *Pba*-infected mice (day 6 PI) (EPs-CM, n=6 independent isolation experiments) and non-infected control mice (EPs-NI, n=3 independent isolation experiments). Exposure to LPS (1 μg/ml) and Poly I:C (5 μg/ml) were used as positive response controls. **e**, Measurement of luciferase activity in IFNβ-reporter BECs exposed to two different doses (2×10^5^ and 2×10^6^) of erythrocytes isolated from the blood of non-infected (NIE) or *Pba*-infected mice (IE), or with EPs obtained from IE-cultures centrifuged at 20 000g. One representative experiment of two performed in duplicate or triplicate. **(F)** Activation of *Ifnβ1* gene expression measured by luciferase activity in IFNβ-reporter BECs treated or not with NH4CL (20 mM) and then exposed to EPs collected from *Pba*-IE cell cultures. One representative experiment of three performed in triplicate is shown. **(G)** *Ifnβ1* gene expression measured by luciferase activity in IFNβ-reporter WT or STING1 KO BECs stimulated with EPs. Results comprise data of two independent experiments done in triplicate. Treatments are compared between the two genotypes by two-way ANOVA using Tukey’s multiple comparisons test. Data are presented as mean values ± SD in **(D**-**G)** and analyzed by one-way ANOVA using Tukey’s multiple comparisons test in **(D**-**F)**.

In BECs cultures, particles isolated from the blood of mice with CM activated IFNβ gene transcription (Fig 4D) and protein expression (fig. S4C), but not those from non-infected mice. Similarly, EPs obtained from IE cultures activated *Ifnβ1* transcription measured by the luciferase activity in IFNβ-reporter BECs (Fig. 4E). In contrast, neither erythrocytes from non-infected mice (NIE) nor erythrocytes from *Pba*-infected mice (IE) induced luciferase activity (Fig. 4E). To determine whether endosomal acidification was required for endothelial response, cells were pretreated with ammonium chloride, which elevates the intralysosomal pH (35). Remarkably, induction of IFNβ in response to EPs was abolished in BECs pre-exposed to ammonium chloride (Fig. 4F). Furthermore, luciferase activity was lower in IFNβ-reporter STING1 KO BECs stimulated with IE-derived EPs (Fig. 4G). This result indicates that *Sting1* deletion in BECs reduces induction of IFNβ by IE-derived EPs. In sum, EPs derived from infected erythrocytes are processed in acidic compartments in BECs, activate STING1 and induce IFNβ.

### Particles emanating from the parasite’s digestive-vacuole induce *Ifnβ1* and *Cxcl10* **in brain endothelial cells**

To identify bioactive factors associated with EPs that trigger type I IFN response in BECs, we established a protocol to fractionate bulk EPs obtained from *P. berghei*-IE cultures on basis of a combination of differential centrifugation, paramagnetic separation and presence of host erythrocytic membrane (Fig. 5A). We found that compared to total EPs, the fraction (Fr1) pulled down at lower centrifugation (5630 g) induced higher *Ifnβ1* transcription measured by luciferase activity (fig. S5A). Therefore, we further fractionated Fr1 using a magnetic LS column to separate a fraction enriched in particles containing iron and hemozoin (Fr2) from the flow through (Fr3) (Fig. 5A). To isolate particles with erythrocytic membrane (TER119^+^), Fr3 was incubated with biotinylated anti-TER119 antibody and streptavidin magnetic beads (Fig. 5A). The retained fraction in immuno-magnetic separation originated the Fr4, and the flow through originated Fr5. Flow cytometry analysis (Fig. 5B and fig. S5B) showed that compared to Fr1, Fr2 and Fr5 are enriched in TER119 negative particles. *Pba*-GFP parasites were used to generate IE cultures, thus, the GFP signal in Fr5 indicate the presence of parasite-derived material. TEM of fixed samples confirmed that Fr2 contained vesicles with hemozoin crystals likely derived from parasite’s digestive vacuoles (Fig. 5C). Among all fractions, the hemozoin-enriched Fr2, was the strongest inducer of luciferase activity in BECs. Compared to total EPs (2×10^6^ EPs), a 10-fold lower amount (2×10^5^ EPs) of Fr2 induced luciferase activity (Fig. 5D). In BECs, Fr2 was also a potent inducer of *Cxcl10* (Fig. 5E) and *Irf7* (fig. S5C) mRNA. Importantly, Fr2-induced expression of these genes was about two-fold lower in STING1 KO BECs than in WT BECs and was abolished in IFNβ KO and in quadruple MyTrMaSt KO BECs (Fig. 5E and fig. S5C). Nevertheless, these KO BECs expressed *Cxcl10* and *Irf7* in response to direct stimulation with IFNβ, which indicates that the IFNβ-IFNAR axis was preserved (fig. S5D). These results indicate that Fr2, the IE-derived particle fraction containing digestive vacuole remnants, strongly induces a type I IFN response in BECs, which is at least partially dependent on STING1.

**Fig. 5.**
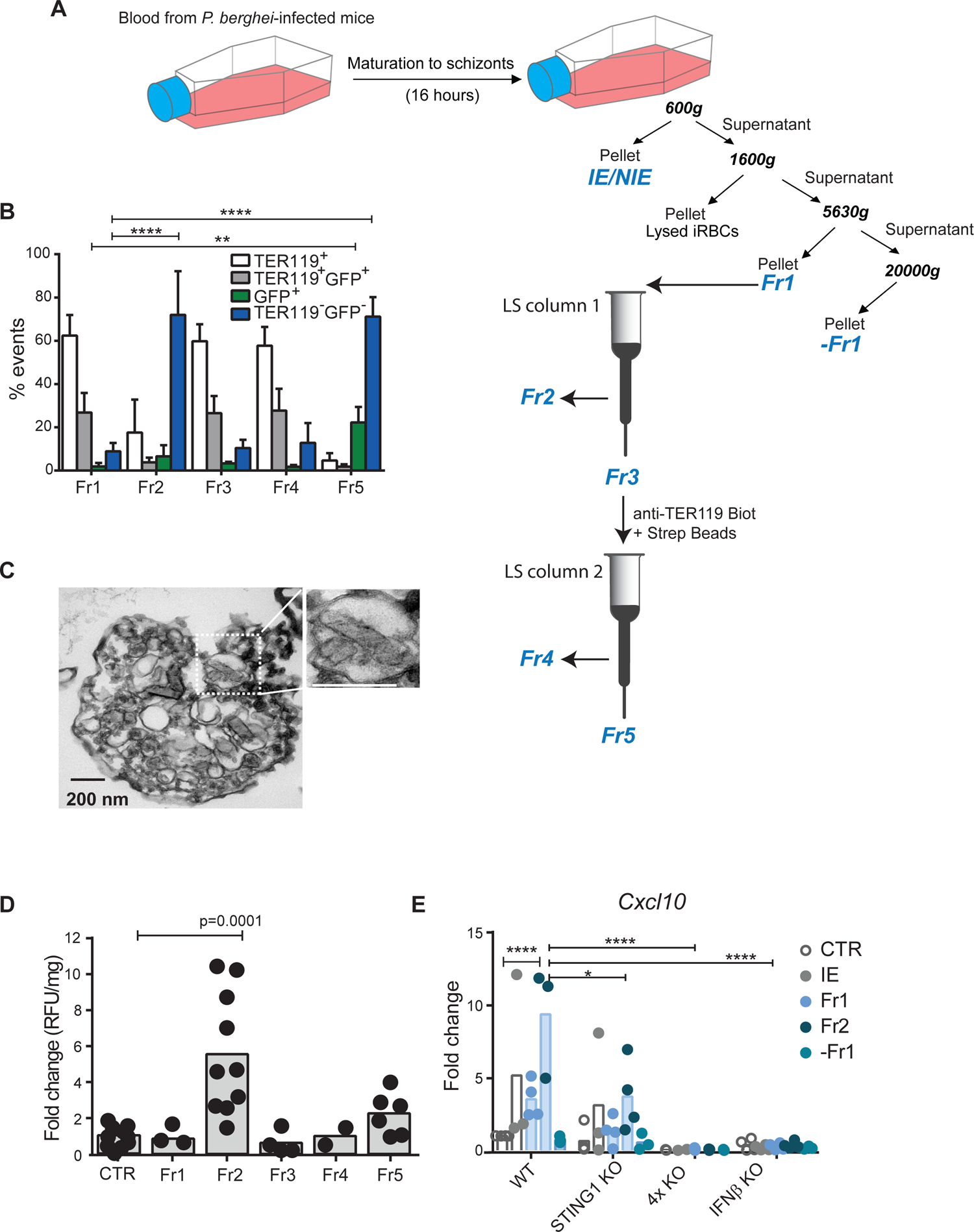
Particles derived from the parasite’s digestive-vacuole activate type I IFN response in brain endothelial cells. **(A)** Experimental workflow for EP fractionation from *Pba*-IE cultures combining multi-step centrifugation with magnetic and immunomagnetic separation to obtain five fractions (Fr1-5). Fr2 is isolated by magnetic field capture of iron-enriched EPs. Fr4 is obtained by immuno-separation of EPs expressing the erythrocyte surface marker Ter119. **(B)** Flow cytometry analysis of EPs staining for TER119 and/or GFP-positive. Data comprising four independent experiments were compared to Fr1 by two-way ANOVA, Dunnett’s multiple comparisons test. ****p<0.0001, **p<0.01. **c**, TEM imaging of Fr2 (fixed sample) showing membrane structures surrounding hemozoin crystals. **(D)** Measurement of luciferase activity in IFNβ-reporter BECs stimulated with each one of the 5 different EPs fractions; at 2×10^5^ EPs/fraction. Data comprise two (Fr4), four (Fr1 and Fr3), six (Fr5) and ten (Fr2) independent isolation experiments and each spot represents a cell culture well. One-way ANOVA, Tukey’s multiple comparisons test. **(E)** *Cxcl10* gene expression in BECs stimulated with blood of *Pba*-infected mice containing 2×10^5^ IE and with 2×10^5^ EPs of different fractions. Data comprise 3-4 experiments of different BECs and EPs isolations and are compared between genotypes by two-way ANOVA using Tukey’s multiple comparisons test, *p<0.05, **** p<0.0001. Each spot represents a cell culture well.

### Labile heme originated by the parasite activates STING1 in brain endothelial cells

We then addressed the role of hemozoin in STING1 activation by inhibiting the hemozoin synthesis pathway in IE cultures with chloroquine, an anti-malarial drug described to increase the levels of labile heme in the parasite (15). Unexpectedly, EPs obtained from chloroquine-treated IE cell cultures induce stronger IFNβ expression than EPs from untreated IE (Fig. 6A). Both hemozoin and labile heme can activate PRRs and contribute to inflammatory storm and BBB dysfunction in CM (14, 16, 17). We stimulated BECs directly with different amounts of heme and hemozoin. While hemozoin was unable to stimulate BECs, 20 μM of heme induced significant luciferase activity in WT but not in STING1 KO IFNβ-reporter BECs (Fig. 6B and fig. S5E). Moreover, heme induced *Cxcl10* gene expression in WT BECs, as previously described (36), but not in STING1 KO BECs (Fig. 6C).

**Fig. 6.**
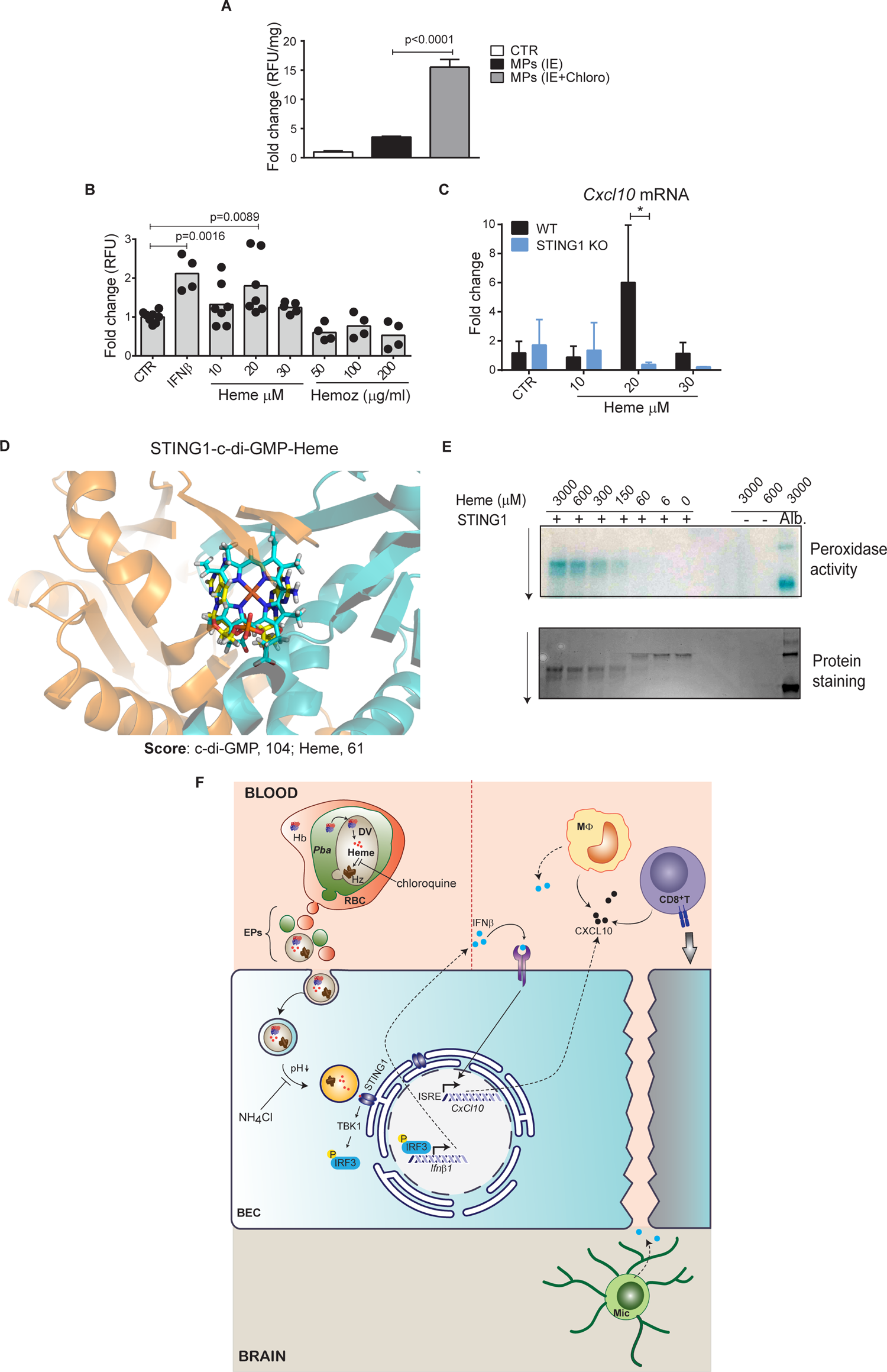
Heme sequestered by the parasite induce *Ifnβ1* and *Cxcl10* gene expression in BECs through STING1 activation. **(A)** Luciferase activity in IFNβ-reporter BECs exposed to 2×10^6^ EPs from *Pba*-IE cultured in the presence or absence of chloroquine (8 μM). One representative experiment of two performed in triplicate is shown. **(B)** Luciferase activity in IFNβ-reporter BECs stimulated with hemin or hemozoin at the indicated concentrations or with IFNβ (1,000 U/ml). Data comprise 2-3 independent experiments and each point represents a cell culture well. **(C)** *Cxcl10* gene expression in WT or STING1 KO BECs stimulated with different concentrations of hemin for 20h. Data comprise two independent experiments. Comparison between genotypes were performed by two-way ANOVA, Tukey’s multiple comparisons test, * p<0.05. **(D)** STING1-Heme molecular docking results. Close-up view of STING1 pocket with superposition of best-scoring docked pose of c-di-GMP and Heme. c-di-GMP and Heme are colored in yellow and cyan sticks, respectively. Molecular docking was performed using GOLD software and ChemPLP as scoring function. **(E)** Detection of peroxidase activity and migration shift assay of STING1 (1μM) incubated with increasing doses of heme. Albumin (5 μM) incubated with heme was used as positive control for peroxidase activity and heme alone was used as negative control. **(F)** Schematic model of STING1 activation in brain endothelium by parasite-derived heme-containing particles and its contribution to the pathogenesis of cerebral malaria. Hb, hemoglobin; DV, digestive vacuole; *Pba*, *Plasmodium berghei*; RBC, red blood cell; BEC, brain endothelial cell; MΦ, monocyte; Mic., microglia; TBK1, TANK binding kinase 1; IRF3; interferon regulatory factor 3; ISRE; interferon-sensitive response element. Data are presented as mean values ± SD in **(A**-**C)** and analyzed by one-way ANOVA using Tukey’s multiple comparisons test in **(A)** and **(B).**

Following these results, we tested a putative interaction of heme with STING1 by performing docking calculations. GOLD software was used to dock heme into the structure of STING1 and search for the optimal pose (orientation, conformation and binding energy) between them. We found that the heme is positioned within the STING1 binding site for its ligand c-di-GMP, mostly overlapping with it (Fig. 6D). This shows that heme can position itself in the pocket in a propitious way for interaction with the STING1. Although well positioned, heme has less affinity to STING1 than c-di-GMP (ChemPLP score of 61 for heme compared to 104 for c-di-GMP). We next tested the ability of STING1 to bind heme by examining the peroxidase-like activity and migration shift of recombinant human STING1 upon incubation with heme. We detected peroxidase activity of STING1-heme complexes at the molar ratio of 1:150, while protein shift was detected at a molar ratio of 1:60 (Fig 6E), values that are similar to those reported for other heme-binding proteins in this assay (37). Taken together, the results indicate that heme-containing EPs derived from IE are able to induce type I IFN responses in BECs in a STING1-dependent manner, possible involving direct heme binding to STING1.

## Discussion

We identified STING1 activation in brain endothelial cells as a molecular pathway that mediates neuroinflammation and brain pathology underlying the pathogenesis of CM. We demonstrated that, early in infection, the brain endothelium uses STING1 to sense *Plasmodium* blood-stage infection and initiate a type I IFN response through activation of *Ifnβ1* and *Cxcl10* gene transcription. This is associated with leukocyte recruitment and subsequent loss of BBB function, a hallmark of CM. We found that extracellular particles derived from IE, containing hemozoin and enriched for labile heme, trigger STING1-dependent IFNβ responses in brain endothelial cells (Fig. 6F).

Our study identifies IFNβ as a critical trigger of IFNAR signaling and a determinant of BBB disruption and lethality in CM. Whole and cell-specific IFNβ-reporter mice enabled us to identify BECs, as well as brain monocytes/microglia, as main producers of IFNβ, and STING1 as the key PRR upstream of IFNβ induction. STING1—a main activator of *Ifnβ1*transcription—also controlled early *Cxcl10* gene induction in the brain of *Pba*-infected mice. CXCL10 is known to promote chemotaxis of CXCR3 expressing CD8^+^ T cells to the central nervous system during *Toxoplasma* infection (28) and has been strongly linked to CM severity both in humans (38, 39) and in rodents (27). We observed that both IFNβ KO and STING1 KO have lower numbers of CD8^+^ T cells and monocytes in the brain infiltrates. Thus, our data support a role for CXCL10 in inducing chemotactic leukocyte migration into the brain.

We provide the first demonstration that activation of STING1 in BECs promotes CM pathology. Infected mice whose brain endothelial/ependymal cells are made deficient in STING1 (Slco1c1CreERT STING1 floxed mice) displayed milder neurological deficits and increased survival. Several studies reported that plasmodial DNA activates STING1 in innate immune cells such as monocytes and macrophages (12, 19, 20). However, we found that specific *Ifnβ1* or *Sting1* deletion in myeloid and microglia cells did not affect CM development in mice. Instead, our data indicate that BECs are the main cell drivers of the STING1-IFNβ axis in the brain during CM.

STING1 is localized in the ER and does not bind DNA directly, but interacts with cyclic dinucleotides, namely cGAMP. cGAMP is produced by cyclic-GMP-AMP synthase (cGAS) upon dsDNA sensing in the cytosol. After translocation into the Golgi apparatus, STING1 and TBK1 phosphorylate IRF3 that promotes transcription of type I IFN genes (40). Interestingly, a population of exosomes isolated from *P. falciparum* (Pf) IE cultures at the ring stage, could transfer Pf-DNA to PBMCs and activate the cGAS-STING1pathway with consequent induction of *Ifn*β1/α gene expression (19). Our study further highlights the likely important role for IE-derived particles in the activation of innate immune mechanisms during malaria. We found that IE-derived particles larger than exosomes and enriched in vesicles containing hemozoin are able to activate *Ifnβ* and *Cxcl10* transcription in BECs. This activation depends on intracellular organelle acidification suggesting a requirement for endosome maturation and/or acidic hydrolase activity (35). IE-derived particles containing hemozoin could represent digestive vacuoles that accumulate heme and/or hemoglobin. Although plasmodial DNA-bound to hemozoin has been shown to activate cyclic GMP-AMP (cGAS) upstream STING1 in macrophages (12), hemozoin itself does not seem to be required for STING1 activation in BECs. In fact, exposure of IE to chloroquine, which reduces hemozoin and increases labile heme bioavailability, enhances IFNβ in BECs. This prompted us to hypothesize that heme could be involved in STING1 activation mediated by IE-derived EPs. Indeed, we observed STING1-dependent up-regulation of *Ifnβ* and *Cxcl10* in the presence of labile heme at concentrations within the ranges found in the plasma of children with CM (41) and known to activate *Cxcl10* transcription in endothelial cells (36). Recently, hemolysis in sickle cell disease and injection of hemin in mice were shown to activate and recruit monocytes to the liver through activation of type I IFN signaling and independently of TLR4 (42). Our docking experiments and protein binding assay indicate that heme can bind STING1 *in vitro*. However, further evidence of direct binding of heme to STING1 in the ER is required to clarify whether heme is a direct or indirect activator of STING1 in BECs. Interestingly, late endosomes tightly contact with the ER (43) indicating that heme-STING1 interaction may occur during the association between acidified mature endosomes containing heme and the ER (Fig. 6f).

Overall, we demonstrate that STING1 in the brain endothelium senses molecular by products of erythrocyte infection by *Plasmodium*, such as labile heme, initiating a type I IFN response mediated by IFNβ with induction of CXCL10 in early phases of infection. This, in turn, triggers a cascade of events that promote brain inflammation, including leukocyte recruitment, and may also affect neuronal responses. This study suggests that type I IFN targeted strategies may minimize brain immunopathology, thus preventing fatal CM and subsequent neurological sequels in survivors.

### Material and Methods

#### Mice and treatments

All procedures involving laboratory mice were in accordance with national (Portaria 1005/92) and European regulations (European Directive 86/609/CEE) on animal experimentation and were approved by the Instituto Gulbenkian de Ciência Ethics Committee and the Direcção-Geral de Alimentação e Veterinária, the national authority for animal welfare.

Mice were housed and bred in the facilities of Instituto Gulbenkian de Ciência. *Myd88^-^*^/-^ (MyD88 KO) mice were originally a gift from Prof. Shizuo Akira (Osaka University). *Mda5^-/-^*mice were obtained from Dr. Maria Mota (IMM, Portugal). Transgenic mice expressing Cre under the GFAP promoter (GFAPCre) in C57BL/6 background were obtained from Dr. Henrique Veiga Fernandes (Champalimaud Foundation, Portugal).

*Ifnβ1^−/−^* (IFNβ KO), *Ifnβ1^+/^*^Δβ-luc^ (IFNβ-reporter mice), *Ifnβ1*^floxβ-luc/floxβ-luc^ (IFNβ ^floxβ-luc/floxβ-luc^), *Trif^-/-^* (TRIF KO), B6(Cg)-*Tmem173^tm1.2Camb^/*J (*Tmem173^−/−^*, STING1 KO), Tg(Slco1c1-icre/ERT2)1Mrks (Slco1c1CreERT2) and *Myd88^−/−^Trif^−/−^Mavs^−/−^Sting1^−/−^* (MyTrMaSt KO) mice were obtained from the TWINCORE, Centre for Experimental and Clinical Infection Research, Hanover, Germany.

C57BL/6J, B6;129-*Mavs^tm1Zjc^/*J (MAVS KO), B6;SJL*-Sting1^tm1.1Camb^*/J (STING1^flox/flox^), B6.Cg-Tg(Tek-cre)1Ywa/J (Tie2Cre), B6J.B6N(Cg)-*Cx3cr1^tm1.1(cre)Jung^*/J (CX3CR1Cre), B6.129P2(Cg)-*Cx3cr1^tm2.1(cre/ERT2)Litt^*/*WganJ* (CX3CR1CreERT2) and B6.129P2-*Lyz2^tm1(cre)Ifo^/J* mice (LysMCre) were originally purchased from The Jackson Laboratory (Bar Harbor, ME) (Table S1). MAVS KO mice were backcrossed to C57BL/6J mice up to 96.82% of recipient genome. IFNβ^floxβ-luc/floxβ-luc^ and STING1^flox/flox^ were crossed with the different Cre lines to generate cell lineage conditional IFNβ-reporter, IFNβ KO mice or STING1 KO mice. Experimental animals carrying one loxP-flanked and one KO allele of *Sting1* were generated by crossing Slco1c1Cre^ERT2^STING1^flox/flox^ or CX3CR1Cre^ERT2^ STING1^flox/flox^ mice with STING1 KO mice. The efficiency of the conditional *Sting1* deletion was expected to be improved by having just one floxed allele. Crossing of STING1^flox/flox^ mice with CX3CR1CreSTING1^+/+^ or Tie2Cre STING1^+/+^ mice generated a deleted loxP-flanking Sting1 allele in the germ line. These mice were then crossed with STING1^flox/flox^ and originated Cre^+^ mice with one deleted *Sting1* allele while another allele remained loxP-flanked and was conditionally deleted. Mice with either a deletion of the floxed allele or a KO allele but also expressing a floxed *Sting1* allele were designated STING1^flox/-^.

Wild type, deleted and floxed alleles of *Ifnβ1* and *Sting1* were identified in experimental animals by genotyping using a 3-primer strategy (Table S2). For induction of *Sting1* deletion, 4-6 week old male and female mice were treated with tamoxifen (2 mg/10 g body weight) in corn oil administered by gavage every other day, totaling five doses. Mice were allowed to recover from tamoxifen treatment for 4 weeks.

For *in vivo* labelling of proliferating cells, mice received four intraperitoneal injections of BrdU (50 mg/kg,) every two hours. Brains were collected for FACS analysis two hours after the last injection and intracardiac perfusion of the mice with PBS. As a positive control for luciferase activity in IFNβ-reporter mice in GFAP^+^ cells, mice received an intraperitoneal injection (i.p.) of 0.4 mg of Poly I:C.

### Parasites and infection

Mice were infected with 10^6^ IE with the following parasite strains: *Plasmodium berghei* ANKA (*Pba*) or *Pba*-GFP (44) to induce CM or with *Pb* NK65, a strain that proliferates in mice but does not induce neurological signs of CM.

Frozen IE stocks were expanded in C57BL/6 mice prior to infection. Parasitemia in mice infected with non-GFP parasites (*Pba* and *Pb* NK65) and GFP-*Pba* was determined by flow cytometry analysis as the percentage of DRAQ5^+^ erythrocytes (blood samples incubated with DRAQ5™, 3 uM) and GFP^+^ erythrocytes, respectively (LSRFortessa™ X-20 cell analyser, BD Biosciences and FACSDiVa software version 6.2).

### Evaluation of CM

#### Neurological score

Wild-type mice developed severe signs of CM such as head deviation, paralysis, and convulsions at day 6–7 post infection (PI) and died within 4–6 h of severe disease development. In different genetic backgrounds death by CM occurred up to 11 days PI (“CM time window”). Mice resistant to CM died around day 20 with hyperparasitemia (∼60%) without signs of severe neurological dysfunction.

Disease severity was evaluated in mice before and after infection following an adapted health score for CM (45). From day 4 PI mice were scored based on eleven parameters: gait/body posture, grooming, motor performance, balance, hindlimb clasping reflex, limb strength, toe pinch, aggressiveness, head deviation, paralysis and convulsions. For each parameter a maximum score of two was given to good response or health condition (normal gait, full body extension, clean and sheen hair, exploring of 3-4 corners in 30 sec, extending front feet on the wall of the cage, normal plantar reaction, active pull away, toe pinch reaction, normal aggressiveness, no convulsions, no paralysis or no head deviation). Clear signs of CM (convulsions, paralysis and head deviation) with no strength, no balance and no toe pinch reaction scored zero. Healthy mice scored 22 and mice dying of CM could score between 3-16 points at day 6 PI.

#### Blood–brain barrier integrity assay

At day 6 PI, when C57BL/6J WT mice show signs of CM, the different mouse groups were killed 1 hour after retro-orbital injection of 100 μl of 2% Evans blue (EB) dye (Sigma) and were perfused intracardially with 15 ml of PBS. Brains were dissected, weighed, and pictured. EB retained in the brain tissue was extracted by immersion in 2 ml formamide (Merck) at 37°C in the dark for 48 h and measured by spectrophotometry at 620 and 740 nm (Thermo Scientific™ Multiskan™ GO Microplate Spectrophotometer). Amounts of EB were calculated using an EB standard and expressed as mg EB/g of brain tissue.

#### Immune cells analysis

Spleen (day 5 PI) and brain (day 6 PI) single cell suspensions of infected and non-infected C57BL/6J and KO mice were immuno-stained for T cell and monocyte/macrophage activation cell surface markers. Brain single-cell suspensions were prepared as described before (25). Briefly, mice were perfused with 20 ml of PBS via the left ventricle of the heart. Brains were removed and homogenized in HBSS containing collagenase VIII (0.2 mg/ml) (Sigma). After incubation for 45 min at 37 °C, the digested tissue was minced, forced through a nylon cell strainer of 100 μm (BD Falcon™) and centrifuged. The pellet was resuspended in a gradient of 30% Percoll (Amersham Biosciences) in PBS and centrifuged at 520 g for 30 min at RT to obtain a pellet with brain mononuclear cells.

Spleen and brain mononuclear cells were incubated with anti-mouse Fc block in FACS buffer (2% FCS and 0.01% NaN3 in PBS) for 30 min at 4 °C to block irrelevant binding of the antibodies to the Fc receptor of myeloid cells. After washing, cells were stained with defined cocktails of fluorochrome-conjugated antibodies (Table S3). For intracellular staining, cells were incubated for 5h in RPMI complete medium containing eBioscience™ Cell Stimulation Cocktail (Invitrogen). Cells were then washed and fixed and permeablized with BD Cytofix/Cytoperm™ (BD) before staining with anti-mouse Granzyme B-efluor 450 and anti-mouse IFNγ-PE (BioLegend). Brain cells isolated from *Pba*-infected mice injected with BrdU were stained using the FITC BD BrdU Flow Kit (BD Pharmingen™) according to the manufacturer’s instructions. Stained cells were resuspended in FACS buffer with 5,000 of 10-µm counting beads (SureCount™ Standards, Bang Laboratories) and analyzed by flow cytometry (LSR Fortessa X20™, BD). Data were analyzed with FlowJo v10 software and cell types identified according to the gate strategy (fig. S6)

### Isolation of extracellular particles (EPs)

All the solutions were filtered (0.22 μm) before use. Blood EPs were obtained from mouse blood collected by cardiac puncture with a syringe containing 100 μl of EDTA, 0.5 M. Plasma was prepared by two successive centrifugations at 1500 g for 15 minutes at 4°C. Harvested plasma was further centrifuged at 20 000g for 25 minutes and the pellet containing EPs was ressuspended in 0.2% BSA in PBS (46).

IE-derived EPs were prepared from the supernatant of IE cultures. Whole blood of *Pba* infected mice (day 5-6 PI, 10-20% parasitaemia) was collected by cardiac puncture with a syringe containing heparin and suspended in 5 ml of RPMI 1640 medium (Biowest), followed by centrifugation at 450*g* for 10 min. The pellet was resuspended in RPMI 1640 medium containing Glutamax and Neomycin (0.05 mg/ml), supplemented with 20% FBS (Biowest) without or with chloroquine, 8 μM, and then incubated in culture flasks overnight in an orbital shaker at 36.5°C and 50 rpm. EPs were isolated adapting a previously described procedure (34) (Fig. 5a). Briefly, the cell suspension was first centrifuged at 600g to remove IE and non-IE and then centrifuged at 1600g for 15min to remove cell debris. The supernatant of this centrifugation was then sequentially centrifuged at 5630g and 20 000g for 20 min (total EPs). The pellet resulting from centrifugation at 5630g was designated fraction 1 (Fr1) and the supernatant was further centrifuged at 20 000g producing a pellet, “–Fr1”. Fr1 was resuspended in PBS and passed through a LS magnetic exclusion column according to the manufacturer’s instructions (MACS Milteny Biotec). Particles retained in this column were designated Fr2. The flow through (Fr3) was centrifuged at 10 000g for 10 min and the obtained pellet incubated with biotinylted anti-mouse TER119 antibody (1 μg/ml) (Invitrogen). The pellet was then washed twice and resuspended in 80 μl of labeling buffer (PBS, 2 mM EDTA) containing 20 μl of Anti-Biotin MicroBeads (Milteny Biotec). After incubation for 15 min at 4°C and washing, the pellet was resuspended in separation buffer (0.5% BSA, 2 mM EDTA in PBS) followed by magnetic separation through a second LS column. Particles retained in this column and flow through were designated Fr4 and Fr5, respectively.

### Characterization of EPs

#### NanoSight

Blood-derived MPs were analyzed for particle concentration and size distribution by the NS300 Nanoparticle Tracking Analysis (NTA) system with a red laser (638 nm) (NanoSight –Malvern Panalytical, United Kingdom). Samples were pre-diluted in PBS to a concentration range of 5×10^7^-1×10^9^ particles/ml and 10-50 particles/frame for optimal NTA analysis. Video acquisitions were performed using camera level 16 and threshold 5. Five videos of 30s each were captured per sample. Analysis of particle concentration per milliliter and size distribution were performed using the NTA software v3.4.

#### Flow cytometry

Blood MPs or IE-derived EPs were pre-incubated with Fc-block and then stained with PE-conjugated anti-mouse Ter119, CD41, CD45 and CD105 antibodies (1:100) (Life Technologies) during 20 min. After staining, the samples were washed in PBS (10 000*g*, 10 min, 4°C) and the pellet was resuspended in PBS and acquired with flow cytometer LSRFortessa™ (BD Biosciences, run with FACSDiVa software version 6.2) using side scatter of blue violet laser (SSC-BV) and FSC-PMT tuned to detect small particles. PBS was used to set background signal and unstained samples used to define PE-negative particles. Total EPs numbers were calculated using 5000 counting beads (10 µm) in each sample.

#### Transmission electron microscopy

For negative staining, 5 µl of the pellet of serum MPs in HEPES buffer (10 mM) was adhered to glow discharged Mesh100 grids coated with 2% formvar in chloroform and carbon for 2 min. Following attachment, samples were washed with distilled H_2_O and stained with 2% uranyl acetate for 2 min.

EPs samples were fixed with 2% formaldehyde and 2.5% glutaraldehyde in 0.1 M phosphate buffer (PB) overnight at 4°C. After washing three times, samples were embedded for support in 2% low melting point agarose. After solidifying, samples were post-fixed with 1% osmium tetroxide in 0.1 M PB for 30 min at 4°C. Samples were then washed in PB and distilled water and pre-stained with 1% tannic acid for 20 min at 4°C and with 0.5% uranyl acetate for 1h at RT in the dark followed by dehydration using increasing concentrations of ethanol. After dehydration, samples were infiltrated and embedded using a graded series of Embed-812 epoxy resin polymerized at 60°C overnight. Ultrathin sections (70 nm) were obtained with a Leica UC7 Ultramicrotome and collected in 2% formvar in chloroform-coated slot grids that were post-stained with 1% uranyl acetate and Reynold’s Lead Citrate for 5 min each. Electron microscopy images were acquired on a FEI Tecnai G2 Spirit BioTWIN transmission electron microscope operating at 120 keV and equipped with an Olympus-SIS Veleta CCD Camera.

### Brain endothelial cell cultures

Mouse brains were removed from the skull and kept in a petri dish with HBSS medium (Biowest) on ice. The brains were transferred to a freezer block covered with gauze and the meninges removed by rolling a cotton swab over each brain. Brain blood vessels were then isolated adapting a previously described procedure (47). Briefly, brains were minced with a blade in 1 ml of HBSS medium and aspirated with a syringe. The tissue was then homogenized by pushing it five times through a 23G needle. This homogenate was mixed with equal volume of cold 30% (wt/vol) dextran (MW ∼60 kDa) (Sigma) in PBS and then centrifuged at 10 000g for 15 min. After centrifugation the myelin top layer was removed and the pellet containing the vessels was suspended in DMEM and passed through a 40-μm cell strainer. Five milliliters of medium was added to wash away other cells and myelin debris. The strainer was back-flushed with 5 ml of MCDB 131 to collect the brain vessels. These were digested by supplementing the medium with 1mg/ml of collagenase type IV (Millipore), 10 μg/ml DNAse (Roche) and 2% FBS, followed by incubation for 2-3 hours at 36.5 °C in an orbital shaker. The digested vessels were then centrifuged and suspended in EGM™-2 Endothelial Cell Growth Medium-2 BulletKit™ (LONZA) supplemented with GlutaMax (ThermoFisher), 10% of FBS and 4 μg/ml puromycin, and cultured in 24 well-plates. Medium was changed for fresh medium without puromycin every four days.

When cultures reached 50-80% confluence (∼14 days, 95% of the cells are CD31^+^ by flow cytometry analysis), cells were exposed for 18-20h to different amounts (2×10^5^-2×10^6^) of IE and NIE, EPs fractions (Total EPs, -Fr1 and Fr1 to Fr5), MPs (present in 600 μl of serum), or different activators of IFNβ signaling (IFNβ, 1000 U/ml; LPS, 1μg/ml; Poly I:C, 1-5 μg/ml). To inhibit endosome acidification, BECs were pre-incubated for 20 min with 20 mM of NH4Cl before adding the stimuli.

Heme was added as ferric chloride heme (hemin, SIGMA) to BECs at a final concentration range of 10-30 μM. A freshly prepared stock solution was obtained by dissolving hemin in NaOH 0.1 M for 1h at 37°C and then adding 1M Tris, pH 7.5 to a final concentration of 0.1 M. HCl was added to the solution to bring the pH back to 7.5.

### Molecular docking

STING1-c-di-GMP crystal structure (PDB ID: 6S26) was retrieved from Protein Data Bank (PDB). The crystal structure was then prepared using the Molecular Operating Environment (MOE v2019.01) structure preparation wizard by stripping out the water molecules. Hydrogen atoms were added, and protonation states were assigned using Protonate-3D tool at pH=7.4 of the MOE software package. The co-crystallized ligand (c-di-GMP) was also prepared using the MOE software package. The EHT10 force field was used in all minimization steps. Again, hydrogen atoms were added using the Protonate-3D tool at pH=7.4.

C-di-GMP was then removed and re-docked into the corresponding STING1 crystal structure. Self-docking was performed in GOLD v5.7 software using ChemPLP as scoring function. The docking binding site was centered at threonine T263 (fig. S5G), a crucial residue for c-di-GMP and STING1 interaction, and a search radius of 15 Å was set. Upon self-docking successfully validated the developed protocol, molecular docking of heme was performed, 500 GA runs. Heme followed the same ligand preparation procedure as c-di-GMP.

### Peroxidase activity and gel migration shift assay

Recombinant human STING1 protein (Gentaur) (0.3 μg, 1μM) in 20 mM Tris-HCl, 0.15 M NaCl, pH=8.0 (Tris buffer) was incubated for 15 min at room temperature (RT) with 3000, 600, 300, 150, 60 and 6 μM of heme. After adding Tris sample buffer (pH=6.8) for native PAGE (no SDS or reducing agents), the samples were loaded on 12% Criterion^TM^ TGX^TM^ Stain-Free^TM^ precast gels for PAGE (Bio-Rad). Heme-STING1 complexes were separated by gel electrophoresis in non-denaturing running buffer. Peroxidase-like activity of heme bound to STING1 was detected on the gel after incubation with TMB Substrate Reagent Set (BD Biosciences). After developing a blue-colored band, the gel was washed with 0.5 M sodium acetate (pH=5) and isopropanol (30%). Migration shift of protein-heme complexes (37) was examined after washing the gel in Milli-Q water and staining with Commassie Brilliant Blue R250 (0.05% in 50% of methanol and 10% of acetic acid in H_2_O) for 2 hours at RT, followed by incubation with destaining solution (10% etanol and 7.5% acetic acid in H_2_O). STING1 without incubation with heme and heme alone were used as negative controls for peroxidase-like activity. Serum bovine albumin (3.3 μg, 5μM) incubated with 600 μM of heme was used as a positive control (48).

### Gene expression

RNA was prepared from total mouse brains of mice and from BECs using Trizol and Cells-to-CT kit (Thermo Fisher Scientific), respectively. One microgram of total brain RNA was converted to cDNA (Transcriptor High Fidelity cDNA Synthesis Kit; Roche). A final volume of 20 μl of the cDNA reaction was diluted 1:3 in RNase-free water to be used in qRT-PCR. *Ifnβ1*, *Cxcl10*, *Irf7*, *Irf1* and *Ifit1* mRNA were quantified using Mm00439552_s1, Mm00445235_m1, Mm00516788_m1, Mm01288580_m1 and Mm00515153_m1 TaqMan Gene Expression Assays from ABI, respectively.

### IFNβ-luciferase reporter assay

Tissues were lysed in Glo Lysis Buffer (Promega) followed by centrifugation at 20,000g for 20 min. BECs were gently washed with PBS before lysis. 50 μl of tissue or cell lysates were mixed with 50µl of Bright-Glo™ Luciferase Assay System (Promega) in a 96-well luminometer plate. The luminescence was immediately measured with GloMax® Explorer (Promega) (excitation, 405nm; emission, 500-550nm; integration time, 5). Protein was quantified by Bradford Assay (BioRad) and luciferase activity expressed as RFU per mg of protein.

### Satistical Analysis

Statistical analysis was performed using Prism v6.01 software (GraphPad Software, Inc.). Comparisons between treated and non-treated conditions used two-tailed unpaired t-test, comparisons between multiple groups used one-way ANOVA, comparisons between treated cells of different genotypes used two-way ANOVA as indicated in figure legends. Bars in figures represent the mean (±SD) and each symbol represents one mouse or well as indicated in the figures. Sample size and number of the independent experiments performed can be found in the figures and figure legends.

## Acknowledgments

We gratefully acknowledge the Electron Microscopy Unit at IGC, in particular Ana Laura Sousa for technical support. We thank M. Soares and S. Gomes for discussions and critical reading of the manuscript. This work was supported by ERA-NET NEURON.

## Supplementary Information

**Table S1.**
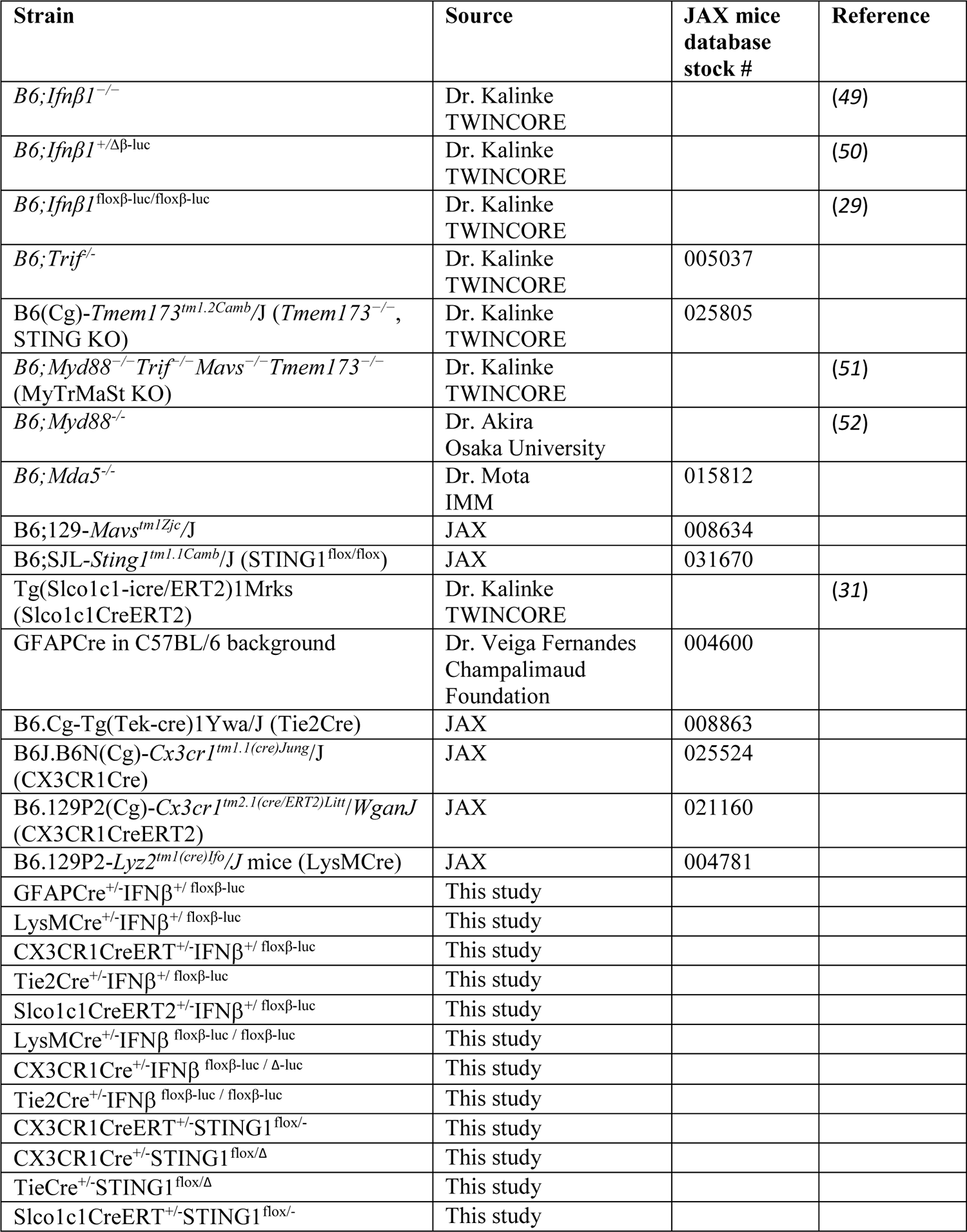
Strains used in this study.

**Table S2.**
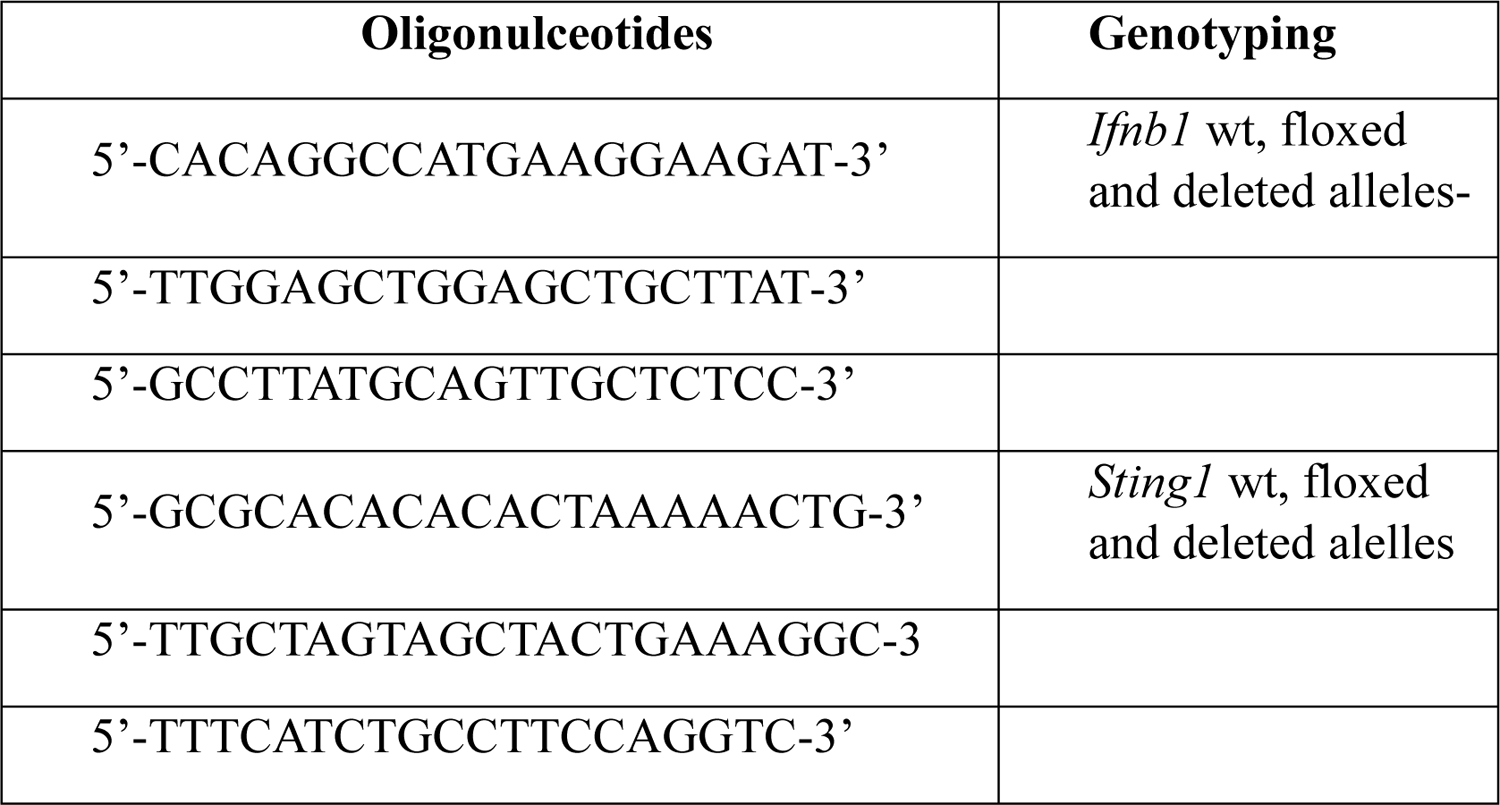
Oligonucleotides used to genotyping *Ifnβ1* and *Sting1* modified alleles.

**Table S3.** Antibodies used for flow cytometry.

| **Antibody** | **Supplier** | **Clone** |
| --- | --- | --- |
| Anti-mouse-Fc-block/CD16/32 | IGC (antibody facility) | 2.4G2 |
| Anti-mouse FITC CD45.2 | IGC (antibody facility) | 104.2 |
| Anti-mouse PE CD45.2 | IGC (antibody facility) | 104.2 |
| Anti-mouse PE CD41 | eBiosciences | MWReg30 |
| Anti-mouse PE CD45 | Life Technologies | 30-F11 |
| Anti-mouse PE CD105 | Life Technologies | MJ7/18 |
| Anti-mouse PE TER119 | Life Technologies | TER119 |
| Anti-mouse efluor 450 CD45.2 | eBiosciences | 104 |
| Anti-mouse biotinylated TCR beta | IGC (antibody facility) | H57-597 |
| Anti-mouse Alexa 648 CD8 | IGC (antibody facility) | YTS169.4 |
| Anti-mouse Pacific Blue CD8 | IGC (antibody facility) | YTS169.4 |
| Anti-mouse biotinylated CD4 | IGC (antibody facility) | GK1.5 |
| Anti-mouse eFluor 450 CD44 | IGC (antibody facility) | IM7 |
| Anti-mouse PE CD69 | IGC (antibody facility) | H1.2F3 |
| Anti-mouse PE CD62L | IGC (antibody facility) | MEL-14 |
| Anti-mouse Alexa 647 CD25 | IGC (antibody facility) | PC61 |
| Anti-mouse biotinylated CD11b/Mac-1 | IGC (antibody facility) | M1/70 |
| Anti-mouse Alexa 488 MHC-I/H-2Kb | IGC (antibody facility) | AF6-88.5 |
| Anti-mouse efluor 450 Granzyme B | BioLegend | N6ZB |
| Anti-mouse PE IFNγ | BioLegend | XMG1.2 |

---

**Fig. S1.**
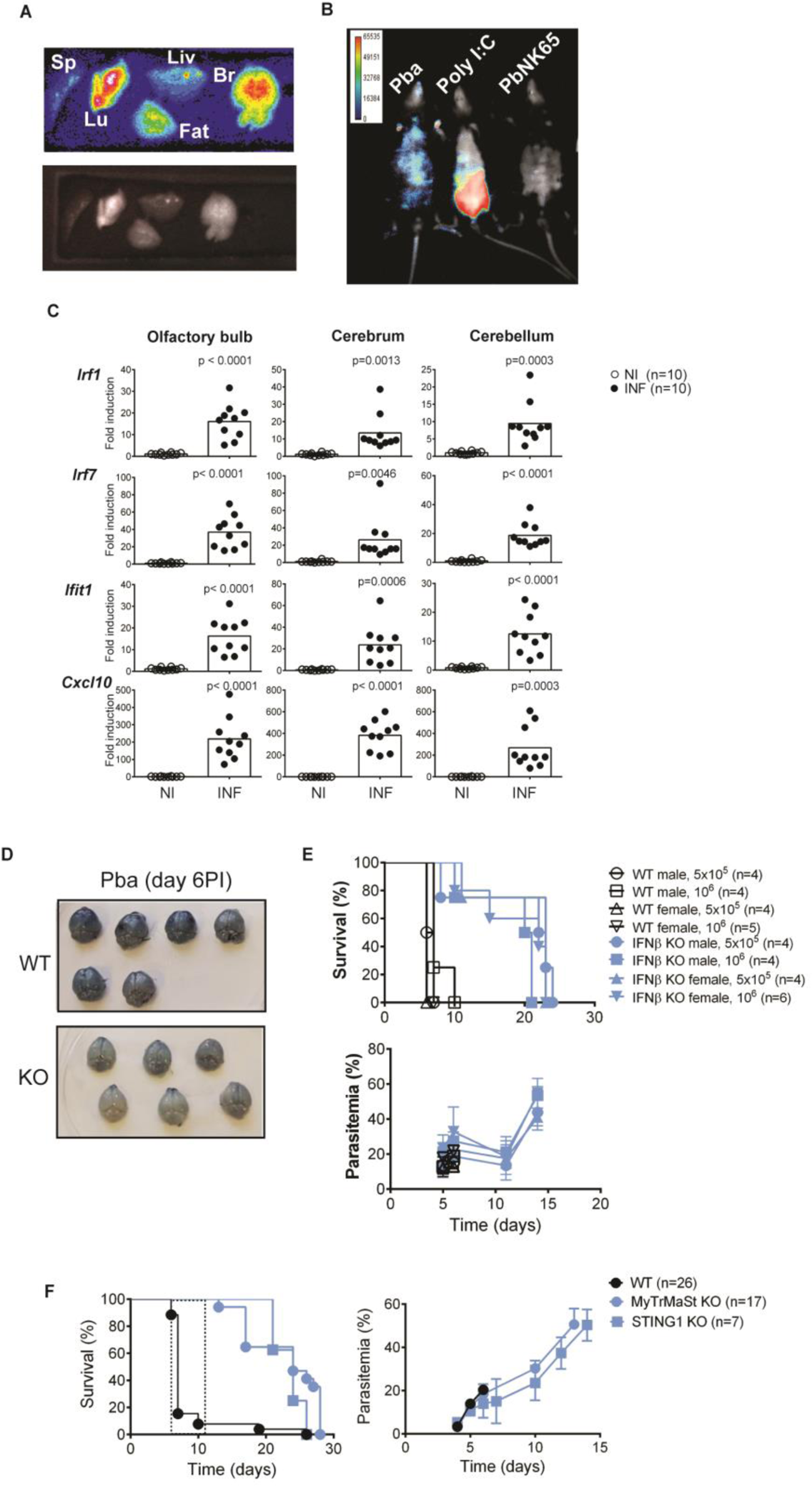
Role of IFNβ signaling in CM development. **(A)** Ex vivo luminescence (upper panel) and corresponding light imaging (lower panel) of spleen (Sp) lung (Lu), liver (Liv), abdominal fat (Fat) and brain (Br) of IFNβ-reporter mice infected with *Pba* (day 5, 20 % parasitaemia). **(B)** *In vivo* luminescence imaging of IFNβ-reporter mice infected with *Pba* (10-20% parasitaemia with CM), injected with a IFNβ inducer (Poly I:C) or with a strain of *P. berghei* that does not induce CM (PbNK65) (day 18, 13% parasitaemia). **(C)** Interferon response signature detected in the designated brain regions was evaluated by measuring interferon-stimulated gene expression by quantitative PCR (qPCR). Data points represent individual mice from two independent infection experiments. Symptomatic *Pba*-infected mice (day 6 PI) were compared to non-infected mice using unpaired, two-tailed *t*-test (p values on the figure). **(D)** Photograph of brains collected from WT and IFNβ KO mice infected with *Pba* (day 6 PI) after Evans blue i.v. and perfusion. **(E)** Survival and parasitemia curves of WT and IFNβ KO male and female mice, infected with the indicated amounts of *Pba*-IE. **(F)** Survival and parasitemia curves of WT, STING1 KO and quadruple KO mice for IFNβ critical components of IFNβ induction (*MyD88*^-/-^, *Trif*^-/-^, *Mavs*^-/-^ and *Sting1*^-/-^) infected with *Pba*. Protection from CM development in quadruple KO mice, is recapitulated in the STING1 KO alone.

**Fig. S2.**
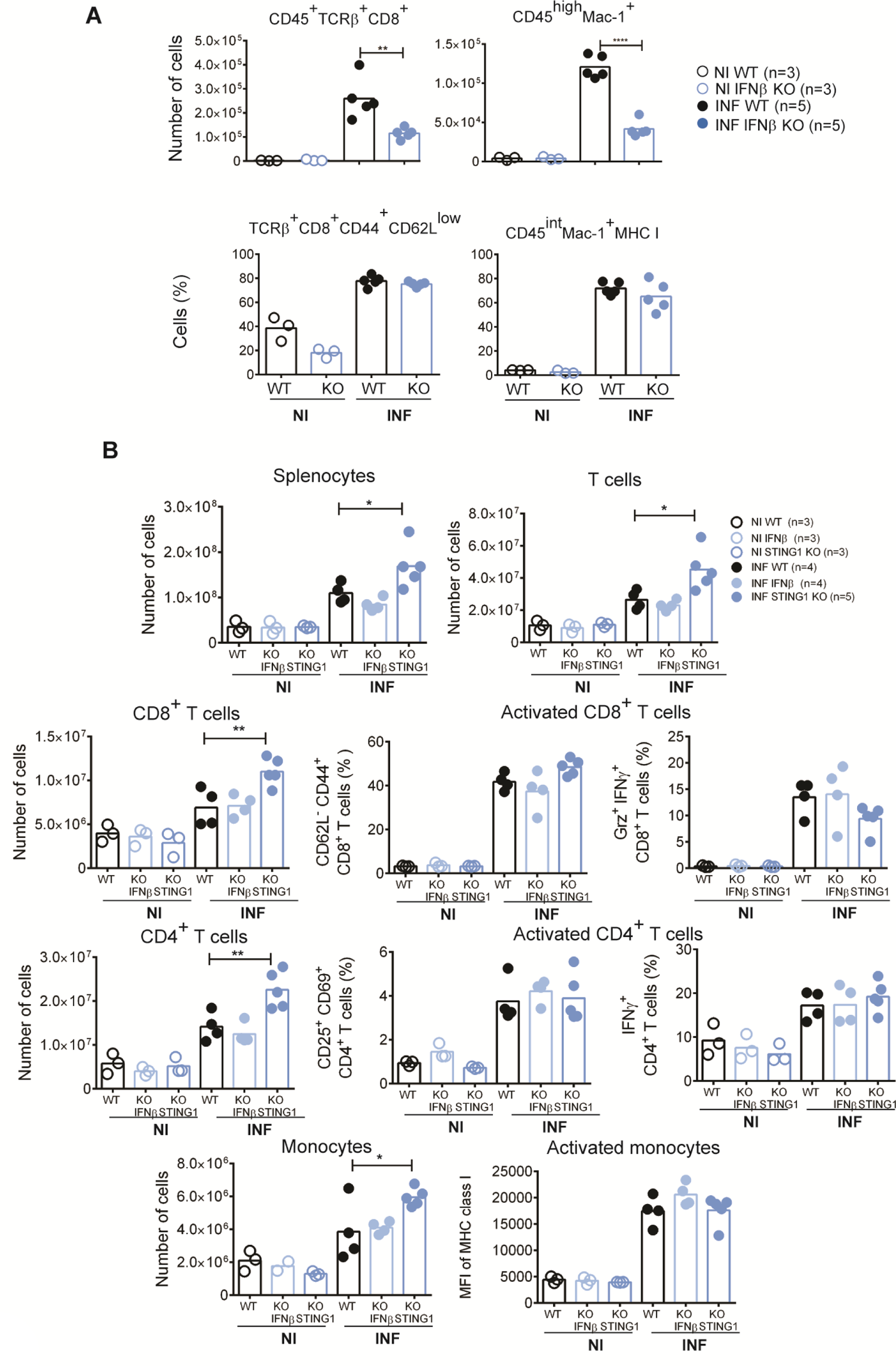
Decreased leukocyte infiltration in the brain is not associated with deficient cell activation. **(A)** Flow cytometry analysis of brain leukocyte infiltration in WT and IFNβ KO mice infected with *Pba*. The number of infiltrating CD8^+^ T cells and monocytes are lower in IFNβ KO while cell activation profiles are maintained. **(B)** Flow cytometry analysis of splenocytes in *Pba*-infected WT, IFNβ KO and STING1 KO mice at day 5 PI. The number of peripheral CD8^+^ T cells, CD4^+^ T cells and monocyte are increased in STING KO mice but the activation profile is not affected. Non-infected (NI) and *Pba*-infected mice of the different genotypes were compared using two-way ANOVA (* p<0.05, ** p<0.01 and **** p<0.0001).

**Fig. S3.**
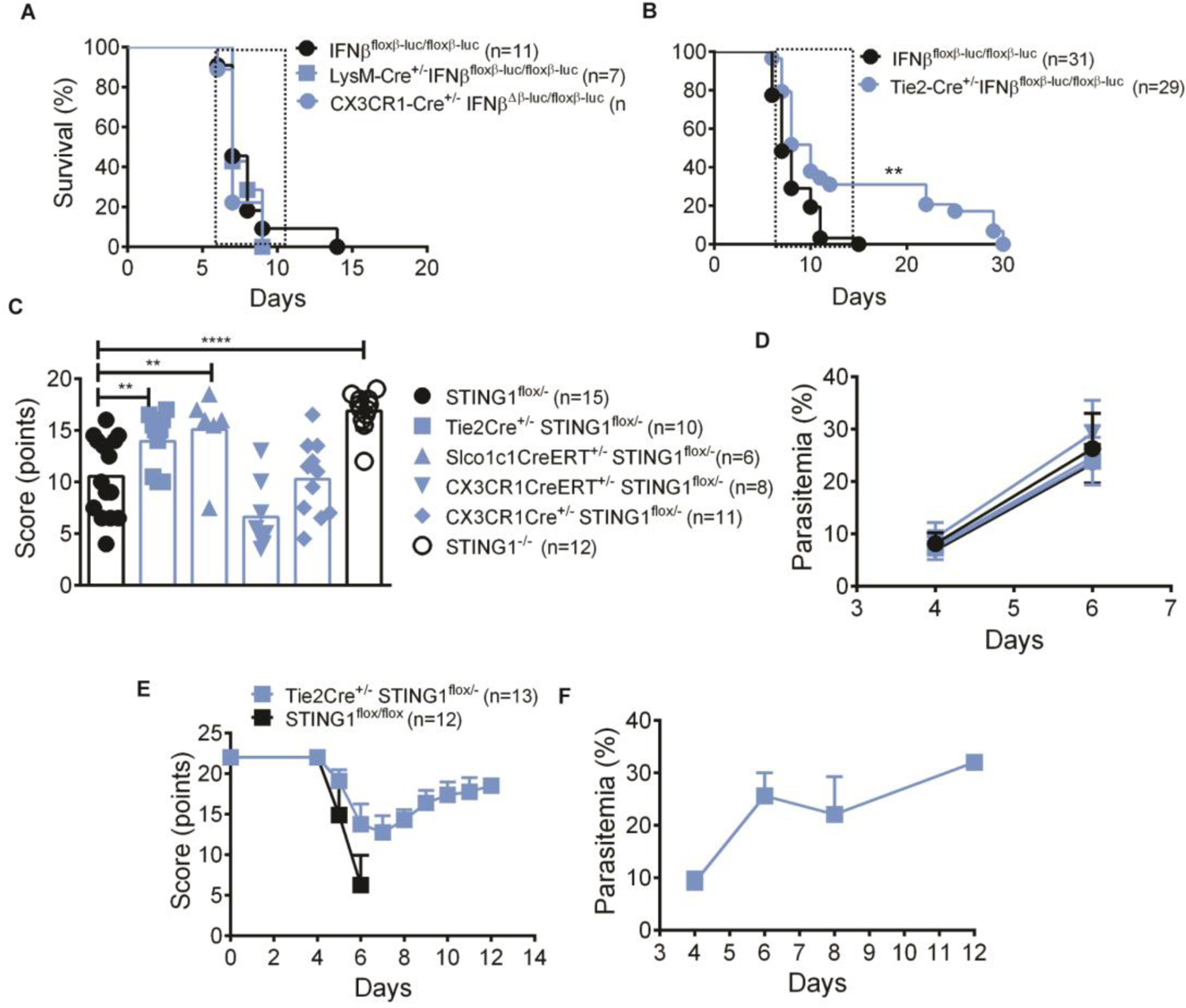
Development of CM in mice with *Ifnβ1* or *Sting1* conditional deletion in myeloid and endothelial compartments. **(A)** Survival of *Pba*-infected mice with conditional IFNβ deletion targeting macrophages (LysM-Cre), microglia (CX3CR1-Cre) and **(B)** endothelial cells (Tie2-Cre). Gehan-Breslow-Wilcoxon test, **p<0.01 **(C)** Health/neurological score of mice with STING-specific deletion in the different cellular compartments. Non-infected animals have a score of 22 that decreases in mice with CM at day 6 PI. Data were compared by unpaired, two-tailed *t*-test. **p<0.01, ***p<0.001 by Mann Whitney test. **(D)** Parasitemia at day 4 and day 6 in conditional STING1 KO mice. **(E)** Kinetics of the health/neurological score, and **f**, parasitemia in Tie2^+^ cell-specific conditional STING1 KO mice.

**Fig. S4.**
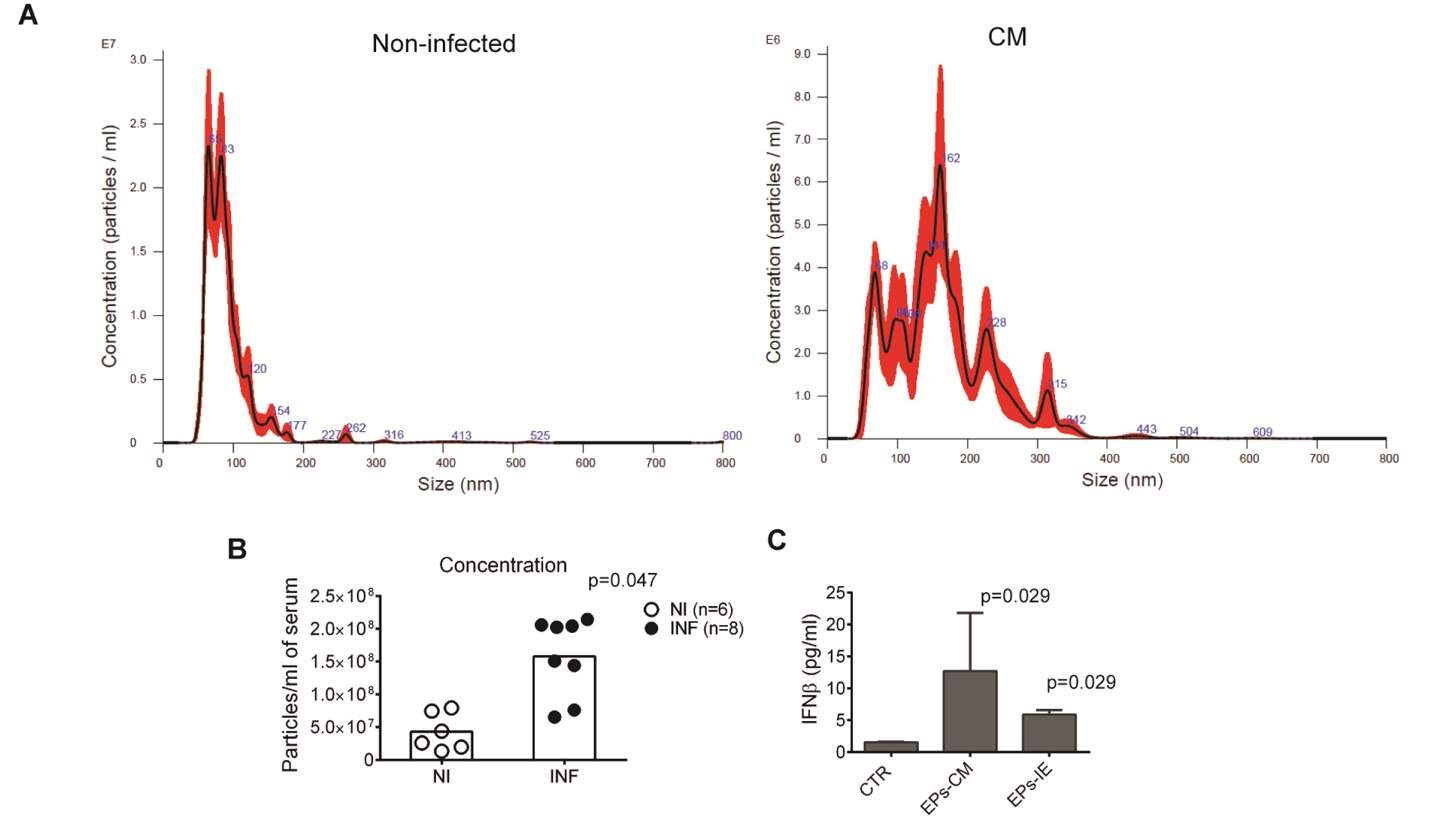
Characterization of extracellular particles obtained from serum of CM affected mice and from *Pba*-IE cultures. **(A)** Representative histogram of the size and concentration of EPs obtained from non-infected and CM wild-type mice. **(B)** NanoSight analysis of concentration of particles within the range of 100.5-265.5 nm in EPs collected from symptomatic *Pba*-infected mice (day 6 PI) and non-infected control mice. **(C)** In vitro IFNβ secretion in brain endothelial cells stimulated with particles obtained from the serum of *Pba*-infected animals (EPs-CM) or *Pba*-IE cultures (EPs-IE), measured by ELISA in cell culture supernatants. Data were compared by unpaired, two-tailed *t*-test.

**Fig. S5.**
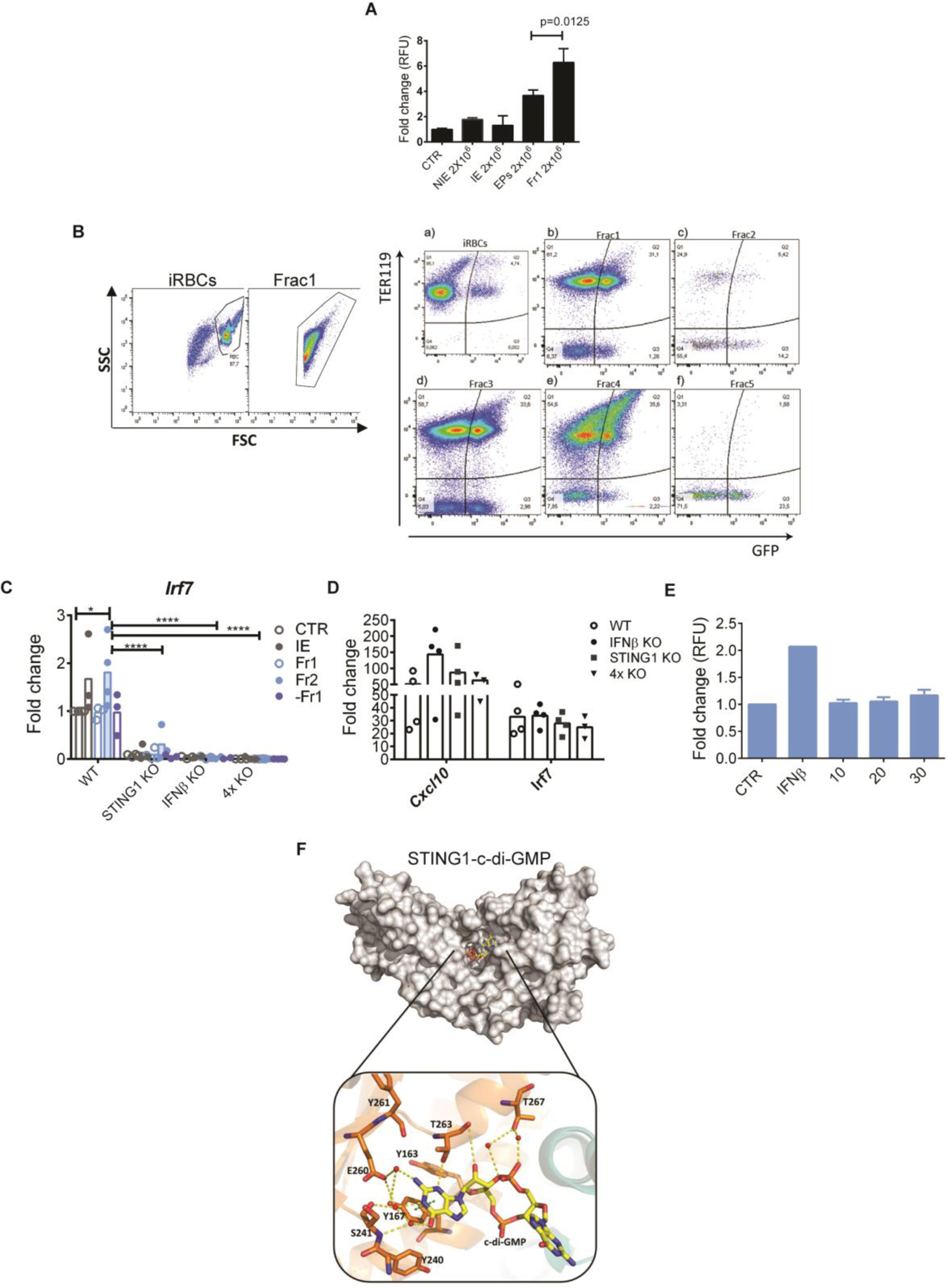
Identification of IFNβ-inducing factors in fractions derived from EPs collected from *Pba*-IE cultures. **(A)** IFNβ gene expression measured by luciferase activity in wild IFNβ-reporter BECs stimulated with 2×10^6^ of total EPs or of Fr1. Results are representative of two independent experiments. One-way ANOVA, Tukey’s multiple comparisons test. **(B)** Flow cytometry profiles of EPs fractions obtained from *PbaGFP*-IE cultures and stained for the erythrocytic marker TER119. **(C)** Induction of *Irf7* expression in BEC of the indicated genotypes upon stimulation with IE and EPs fractions. 4XKO refers to quadruple KO (*Myd88*^-/-^, *Trif*^-/-^, *Mavs*^-/-^ and *Sting1*^-/-^). *Irf7* expression is induced by Fr2 in WT cells but is ablated in absence of endogenous IFNβ expression. Data comprise 3-4 experiments. Comparison between genotypes were performed by two-way ANOVA using Tukey’s multiple comparisons test, *p<0.05, **** p<0.0001. Each spot represents a cell culture well. **(D)** Induction of *Cxcl10* and *Irf7* expression in BEC of indicated genotypes stimulated with IFNβ (1000 U/ml) showing response to IFNAR signaling in the different genotypes. **(E)** Luciferase activity of STING1 KO IFNβ-reporter BECs exposed to different concentrations of hemin for 20 hours. Data comprise two different experiments. **(F)** Surface view of c-di-GMP pocket. Close-up view of specific recognition of c-di-GMP by STING1 and detailed interactions between ribose-phosphate of GMP and water, as well as c-di-GMP and T263 from STING1 monomer. Hydrogen bonds between c-di-GMP and water are shown as yellow dashed lines and red spheres, respectively. π-π interaction of c-di-GMP and Y167 is shown as a green dashed line. Residues from STING1 monomer A that interact with c-di-GMP are shown as orange sticks. c-di-GMP is shown as yellow sticks.

**Fig. S6.**
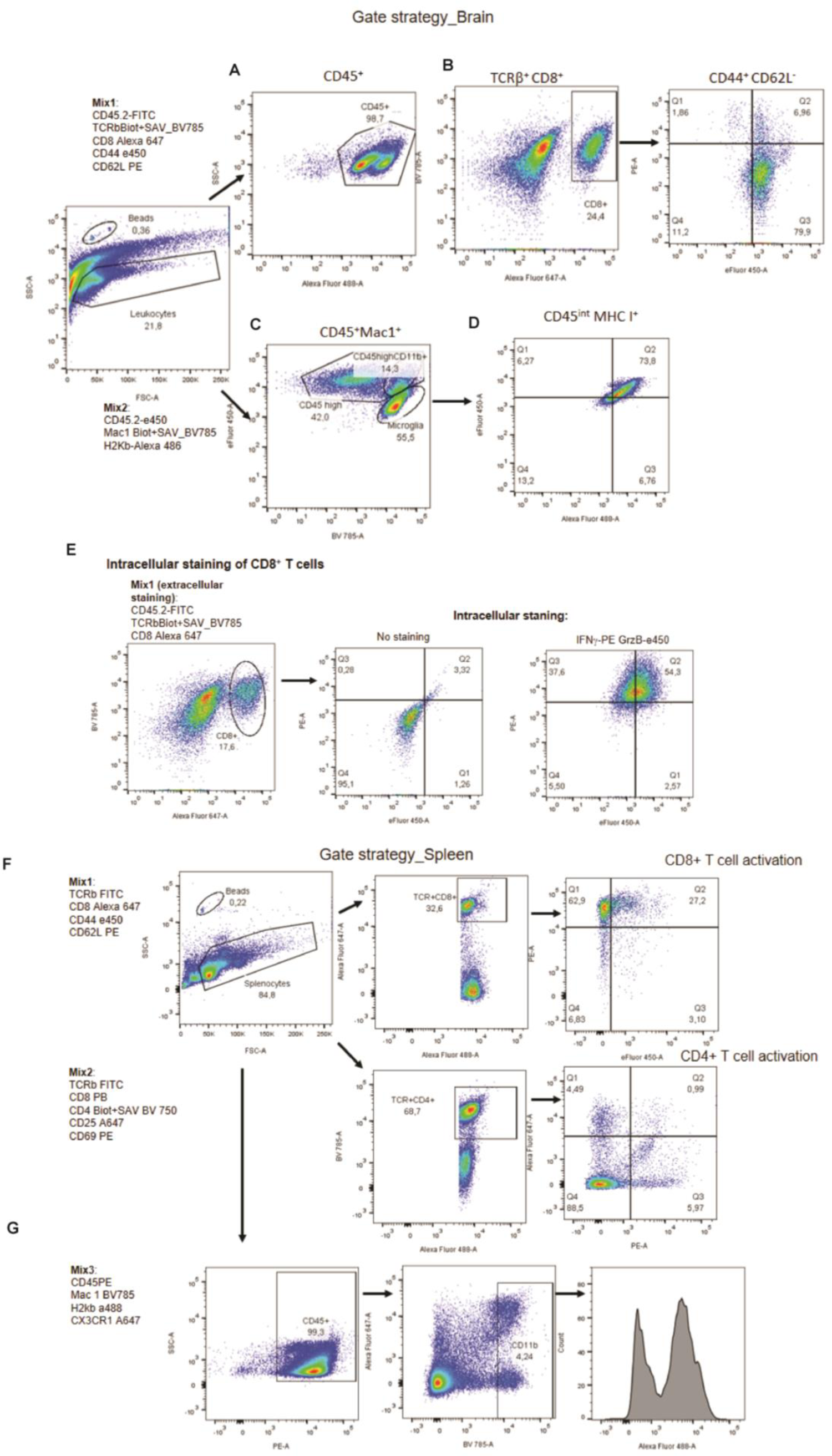
Flow cytometry gating strategy for the identification of immune cells. Brain cells were initially gated on forward scatter (FSC)-A vs side scatter (SSC)-A and then. **(A)** gated as CD45^+^ cells and identified as TCRβ^+^CD8^+^. **(B)** TCRβ^+^CD8^+^ gated cells were examined using cell surface markers for activation (CD44 and CD62L). **(C)** Monocytes and microglia, distinguished based on CD45 expression and Mac-1, were characterized as CD45^high^ Mac-1^+^ and CD45^int^ Mac-1^+^, respectively. **(D)** Within microglial cell population, cellular activation was examined based on MHC class I expression. **(E)** Activation of CD8^+^ T cells was further studied using positivity for IFNγ and Granzyme B. **(F)** Analysis of cell activation profile in splenic CD8^+^ and CD4^+^ T cells and **(G)** monocytes (Mac-1^+^) of experimental mice.

